# A decoy heterotrimeric Gα protein has substantially reduced nucleotide binding but retains nucleotide-independent interactions with its cognate RGS protein and Gβγ dimer

**DOI:** 10.1101/795088

**Authors:** Fei Lou, Tigran M. Abramyan, Haiyan Jia, Alexander Tropsha, Alan M. Jones

## Abstract

Plants uniquely have a family of proteins called extra-large G proteins (XLG) that share homology in their C-terminal half with the canonical Gα subunits; we carefully detail here that Arabidopsis XLG2 lacks critical residues requisite for nucleotide binding and hydrolysis which is consistent with our quantitative analyses. Based on microscale thermophoresis, Arabidopsis XLG2 binds GTPγS with an affinity 100-1000 times lower than that to canonical Gα subunits. This means that given the concentration range of guanine nucleotide in plant cells, XLG2 is not likely bound by GTP *in vivo*. Homology modeling and molecular dynamics simulations provide a plausible mechanism for the poor nucleotide binding affinity of XLG2. Simulations indicate substantially stronger salt bridge networks formed by several key amino-acid residues of AtGPA1 which are either misplaced or missing in XLG2. These residues in AtGPA1 not only maintain the overall shape and integrity of the apoprotein cavity but also increase the frequency of favorable nucleotide-protein interactions in the nucleotide-bound state. Despite this loss of nucleotide dependency, XLG2 binds the RGS domain of AtRGS1 with an affinity similar to the Arabidopsis AtGPA1 in its apo-state and about 2 times lower than AtGPA1 in its transition state. In addition, XLG2 binds the Gβγ dimer with an affinity similar to that of AtGPA1. XLG2 likely acts as a dominant negative Gα protein to block G protein signaling. We propose that XLG2, independent of guanine nucleotide binding, regulates the active state of the canonical G protein pathway directly by sequestering Gβγ and indirectly by promoting heterodimer formation.

## INTRODUCTION

The canonical heterotrimeric guanosine nucleotide-binding protein complex, consisting of Gα, Gβ and Gγ subunits, serves as a molecular on-off switch in the cell. The inactive or “off–state” form consists of the guanosine diphosphate (GDP) bound to the Gα subunit in complex with the Gβγ dimer. For the active or “on–state”, exchange of GDP for GTP in Gα, either spontaneously or catalyzed by a guanine nucleotide exchange factor, changes the Gα conformation leading to dissociation, partly or entirely (1), from the Gβγ dimer and thus enabling both Gα and Gβγ to propagate signaling to downstream components (2–4). Signaling is terminated when the Gα subunit hydrolyzes GTP thus returning to the inactive GDP-bound state. The rate of GTP hydrolysis is an intrinsic property of each Gα subunit but it can be accelerated by Regulator of G protein Signaling (RGS) proteins (5, 6). The Gα structure required for nucleotide binding and hydrolysis and for interaction with Gα-Gβγ and Gα-RGS interactions are well understood (7, 8).

In humans, there are multiple genes encoding G protein subunits resulting in 23 Gα, 5 Gβ and 12 Gγ subunits. The Gα subunits are divided into four subclasses (Gs, Gi, Gq and G12/13) based on function and sequence similarity. However, in Arabidopsis, there is only one canonical Gα (AtGPA1) which approximates the sequence of the ancestral Gα subunit that evolved into these four animal Gα subclasses (9). AtGPA1 has a near identical structure to that of human Giα1 (10). In addition to the canonical Gα subunit AtGPA1, the Arabidopsis genome encodes three atypical Extra-large G proteins (XLG1, XLG2, and XLG3) (11). The other components of the Arabidopsis G protein core are a Gβ subunit (AGB1) (12), one of three Gγ subunits (AGG1, AGG2, and AGG3) (13), and one receptor-like RGS protein (AtRGS1) (14).

The presence of these atypical G proteins makes G protein signaling in plants unique and paradoxical (11, 15, 16). Specifically, the N-terminal half of XLG proteins lacks homology to any characterized domain but contains a putative nuclear localization signal and a cysteine-rich region while the C-terminal half of XLG proteins shares homology (i.e. evolutionary history, (16)) with canonical Gα subunits (~30% identity). However, there is controversy to what extent that these atypical Gα homologs bind and hydrolyze nucleotides and interact with AtRGS1 and AGB1 (17, 18).

For canonical Gα subunits, there are three major conformational changes between the GDP and GTP-bound states of the protein located in what are called Switch I, II and III. Switch I and Switch II directly contact the bound guanine nucleotide and include residues critical for catalyzing GTP hydrolysis, while Switch III contacts Switch II when in the activated conformation (19). These switches are linked between the nucleotide-binding domain and the RGS binding domain and are represented by five conserved sequence motifs named G1 to G5 (20). The G1–G3 boxes provide critical contacts for the β and γ phosphates of the guanine nucleotide and are essential for the coordination of Mg^2+^. The G4 and G5 loops are involved primarily in binding the guanine ring. The G2 and G3 boxes overlap with Switches I and II that are also the key Gβγ binding sites. The RGS domain directly binds to the three switch regions and stabilizes them in a transition state conformation.

It is paramount to resolve unequivocally if XLG proteins bind guanine nucleotide and relevant signaling elements such as RGS and Gβγ to elucidate its atypical mechanism. Here, we combine structure-based and physicochemical experimental methods along with molecular simulations to analyze the binding of XLG2 with both the nucleotide and with a candidate binding partner, AtRGS1/Gβγ dimer. We describe for the first time in great detail the structural issues that should raise concern among those who claim that XLG proteins are nucleotide-dependent switches. In fact, we show that XLG2 binds nucleotide so poorly that it is essentially nucleotide free in the cell, yet despite its nucleotide-free, “empty” state, XLG2 interacts with its partners AtRGS1 and AGB1 with an affinity similar to AtGPA1 in its transition state. We used molecular dynamic simulations to explain how this binding is disrupted and how these protein-protein interactions are maintained.

## RESULTS AND DISCUSSION

### XLG proteins lack critical residues for coordination of the γ and β phosphates on the guanine nucleotide

A multiple sequence alignment of XLGs Gα domain, AtGPA1, and human Giα1 is shown in Fig. 1 with the G1-G5 motifs and switches I-III regions highlighted (noted as SwI-III). To compare the protein structures between XLGs and canonical Gα subunits, we created high-quality models of the Gα homology domains of XLG2 using MODELLER and the aligned sequences shown in Fig. 1. The human RGS4 and Giα1 transition state (Ligand: AlF_4_ and GDP) complex (PDB [1AGR]) was used as template to generate the models of XLG2. Models were created using the *Automodel* script based on the template of human Giα1 in complex with AlF_4_ and GDP (PDB [1AGR]). For evaluation and selection of the “best” model, we calculated the objective function (molpdf) DOPE score, GA341 assessment score between the model and the template (Fig. S1). The final model (XLG2-1) was selected given the lowest average value of the molpdf and the DOPE assessment scores.

**Figure 1.**
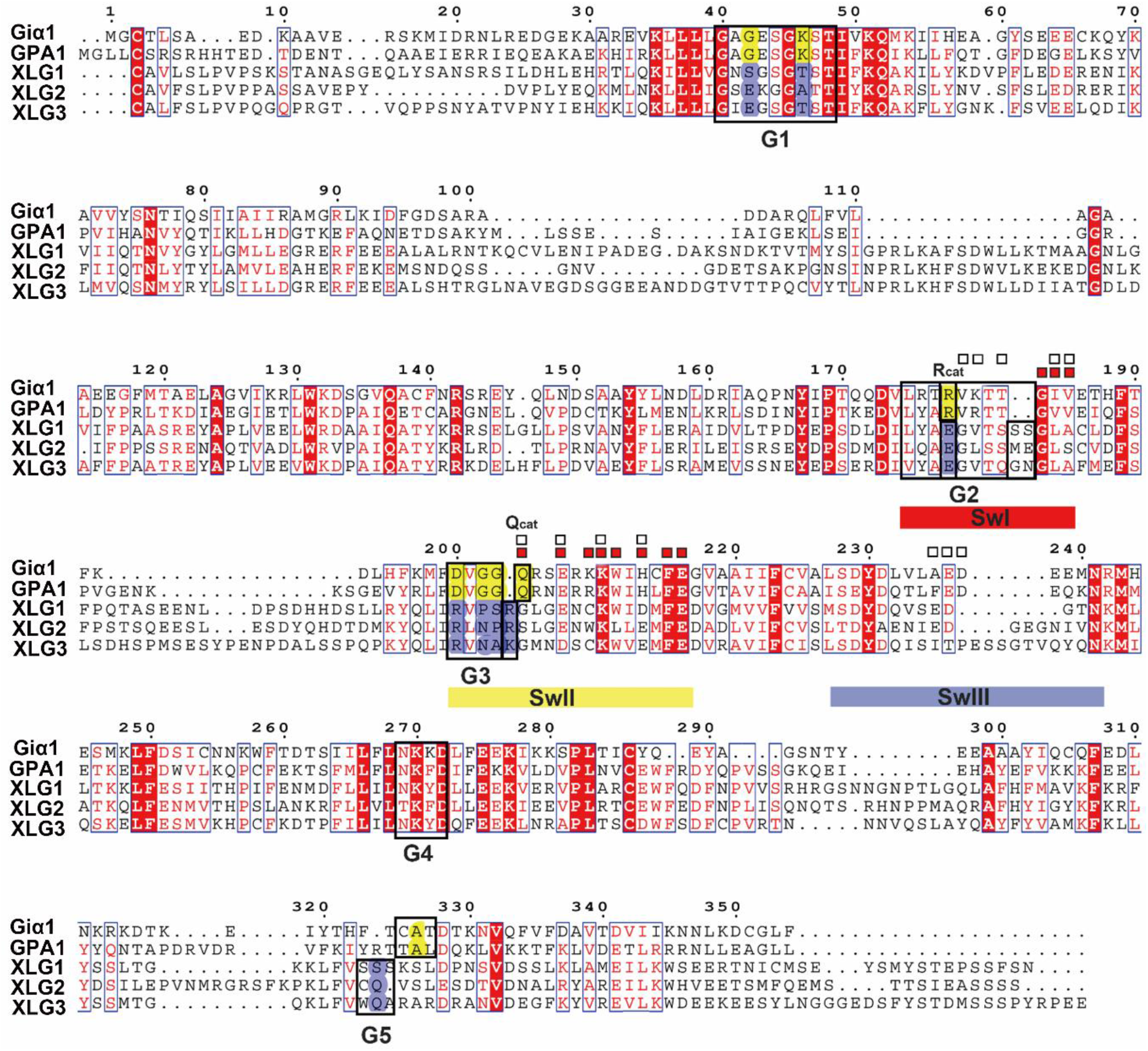
Alignment between human Giα1 Arabidopsis AtGPA1 and the C terminal G alpha domain of the three XLGs The G1-G5 motifs are shown in black boxes. The switches I-III regions (SwI-III) are highlighted (SwI in red, SwII in yellow and SwIII in blue). A percentage of equivalent residues is calculated per columns, considering physico-chemical properties. Blue boxes highlight residues with the same physico-chemical properties and red solid highlighting means the same residues. The contact residues to RGS protein are labeled with white boxes □ and the contact residues with Gβ are labeled with red boxes 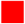. The residues which are conserved in human Giα1 and AtGPA1 for GTP/GDP binding and hydrolysis but are missing in XLGs are highlighted with yellow and blue respectively. The residues essential for the catalysis of the nucleotide are highlighted as Rcat and Qcat. (The C domain of XLGs start with first C residues in the paper, C436 in XLG1, C435 in XLG2 and C396 in XLG3).

As shown in Fig. S2, human Giα1 and Arabidopsis GPA1 have two domains: a Ras-like domain and an all-helical domain. Animal Gα subunits and AtGPA1 are extremely similar in structure (RMSD= 1.8 Ȧ (10)). The Ras-like domain is essential for the nucleotide and RGS proteins binding which contains the five guanine nucleotide binding motifs (G1-G5) and three flexible switch regions (SwI-III) (Fig. S2A). The all-helical domain is important for the intrinsic nucleotide exchange rate (21, 22). XLG2-1 shares a similar overall 3D structure with human Giα1 and AtGPA1 even though the sequence identity is ~ 30%. XLG2-1 contains a globally similar Ras-like domain and α helix domain. The three switch regions and the G1-G5 boxes are highlighted (Fig. S2B). The RMSD between Giα1 and XLG2-1 is 0.67 Ȧ. However, despite similar global structure between XLG2-1 and human Giα1, many of the conserved motifs which are essential for nucleotide binding and hydrolysis are missing, including key residues within the G1, G3 and G5 motifs for nucleotide binding and some dominant residues in the P loop, Switch I and Switch II for coordinating water and Mg^2+^ to catalyze GTP hydrolysis (Fig. 1 and Fig. 2B). These critical differences between the canonical Gα and the XLG Gα domain are described in detail next.

**Figure 2.**
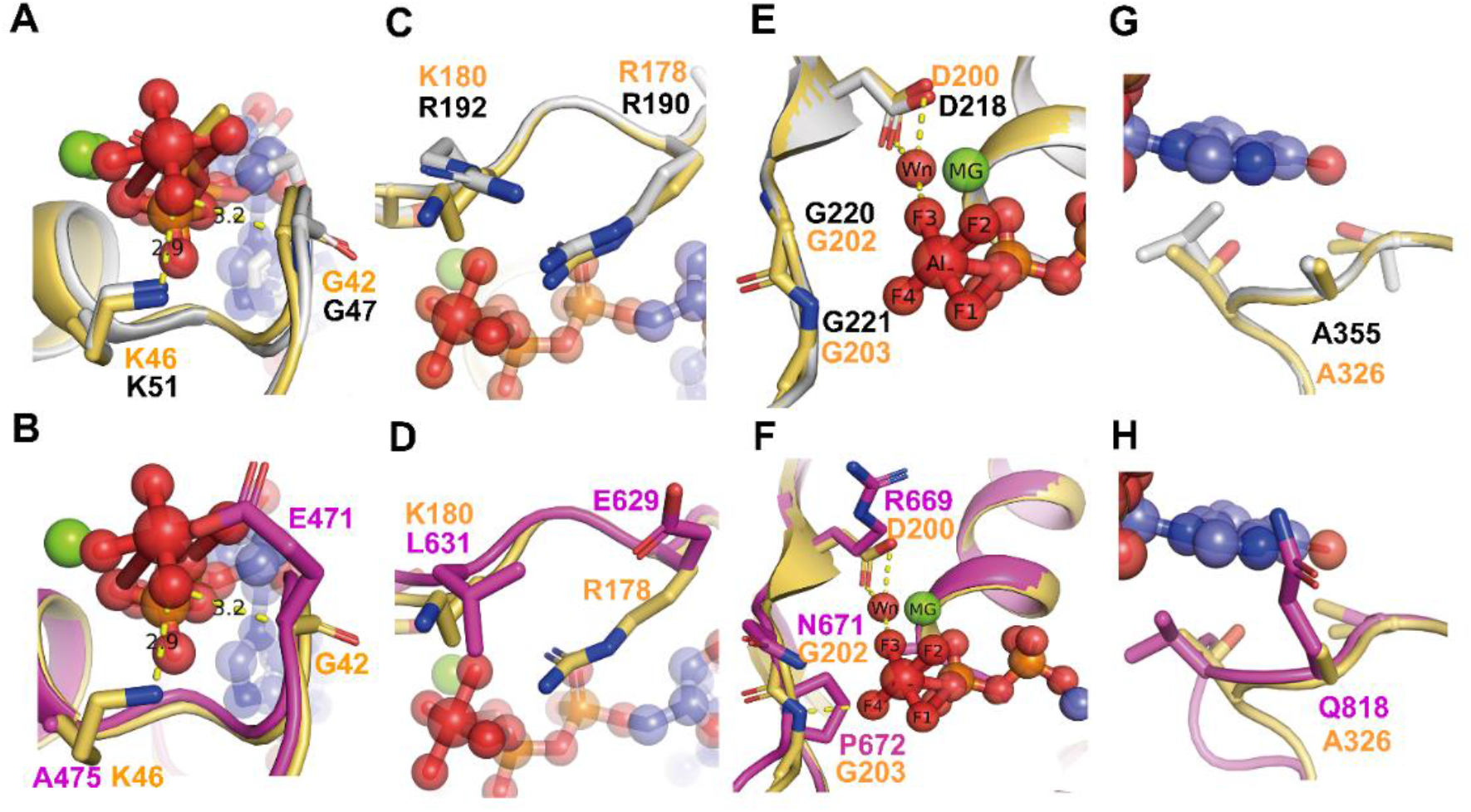
Comparison of the G motifs of AtGPA1 PDB [2XTZ] and the XLG2 G alpha domain model with human Giα1 (PDB [1AGR]) Grey: AtGPA1, Magenta: XLG2-1 model, Light orange: Giα1. The substrate GDP and AlF_4_ are shown as sticks and spheres. Mg^+2^ is shown as green sphere. Wn: nucleophilic water. The main different residues in G1 motif of XLG2 compared with Giα1 and AtGPA1 are shown as sticks **(A, B)**. Both AtGPA1 and Giα1 have the same G and K residues in G1 motif (A). the G42 and K46 in Giα1 were replaced by E471 and Al475 in the counterpart position of XLG2 **(B)**. The main different residues in G2 motif of XLG2 compared with Giα1 and AtGPA1 are shown as sticks **(C, D)**. Both AtGPA1 and Giα1 have the same R residues (known as arginine finger) and similar charged K180 and R192 in G2 motif **(C)**. But in XLG2 no arginine finger exists rather a Glu is at this position. Also, the charged K or R was replaced by a L **(D)**. The main different residues in G3 motif of XLG2 compared with Giα1 and AtGPA1 are shown as sticks E, F. Both AtGPA1 and Giα1 have the same DVGG residues in G3 motif **(E)**. But in XLG2 the DVGG was replaced by R669/N671/P672 relatively (**F**). The main different residues in G5 motif of XLG2 compared with Giα1 and AtGPA1 are shown as sticks in G and H. Both AtGPA1 and Giα1 have the same A residues in G5 motif **(G)**. While in XLG2 the conserved A was replaced by Q818 **(H)**.

The highly conserved G1 motif is a phosphate-binding region containing a flexible structure designated “P-loop” (23). The G1 motif has a consensus sequence of GXXXXGKS/T for the heterotrimeric Gα subunits (7). The P-loop envelopes the phosphates allowing the main chain and side-chain nitrogen atoms to interact tightly with the negatively-charged phosphates (Fig 2 A, B). In animal Gα subunits as well as in AtGPA1, the sequence in the P loop and G1 motif are invariantly set to “GAGESGKS” (Fig 1, see G1 box). However, in XLG2-1, the G42 residue of Giα1 in the P loop is replaced by E471 and the K46 residue of Giα1 is replaced by A475, respectively (Fig 1 and Fig 2B). The G42 residue of Giα1 or G47 residue of AtGPA1 in the P loop play a dominant role in binding the substrate with the main chain forms hydrogen bond with the γ phosphate oxygen atom (23). This G residue is shown in Fig 2A and B. More importantly, only a G residue side chain is small enough to avoid steric clash with the nucleotide and mutation of the corresponding P-loop residue in Giα1, G42 to V, also drastically reduces its GTP hydrolysis activity (24–26). Structural studies of G42V mutant in Giα1 suggest that the introduced valine side chain sterically prevents appropriate positioning of Q204 which coordinates a nucleophilic water molecule during GTP hydrolysis and steric pressure will induce the reconfiguration of switch II (6, 25, 26). Thus, we assume that the substitution of the large side chain of E471 in XLG2-1 reduces GTPase activity (Fig 2B).

Additional differences were found with the P loop of the XLG proteins. The lysine (K46 of Giα1 and K51 of AtGPA1) residue in the G1 motif directly interacts with the β- and γ-phosphate oxygens of the GTP and thus is crucial for the required free energy change (6) (Fig. 2A). Given that there are two dominant residues mutations in the nucleotide pocket of XLG2 (G42 in Giα1 to E471 and K46 to A475) (Fig. 3B), we hypothesize that XLG2 binds the nucleotide with a reduced affinity *in vitro* and that XLG2 is nucleotide free *in vivo*.

**Figure 3.**
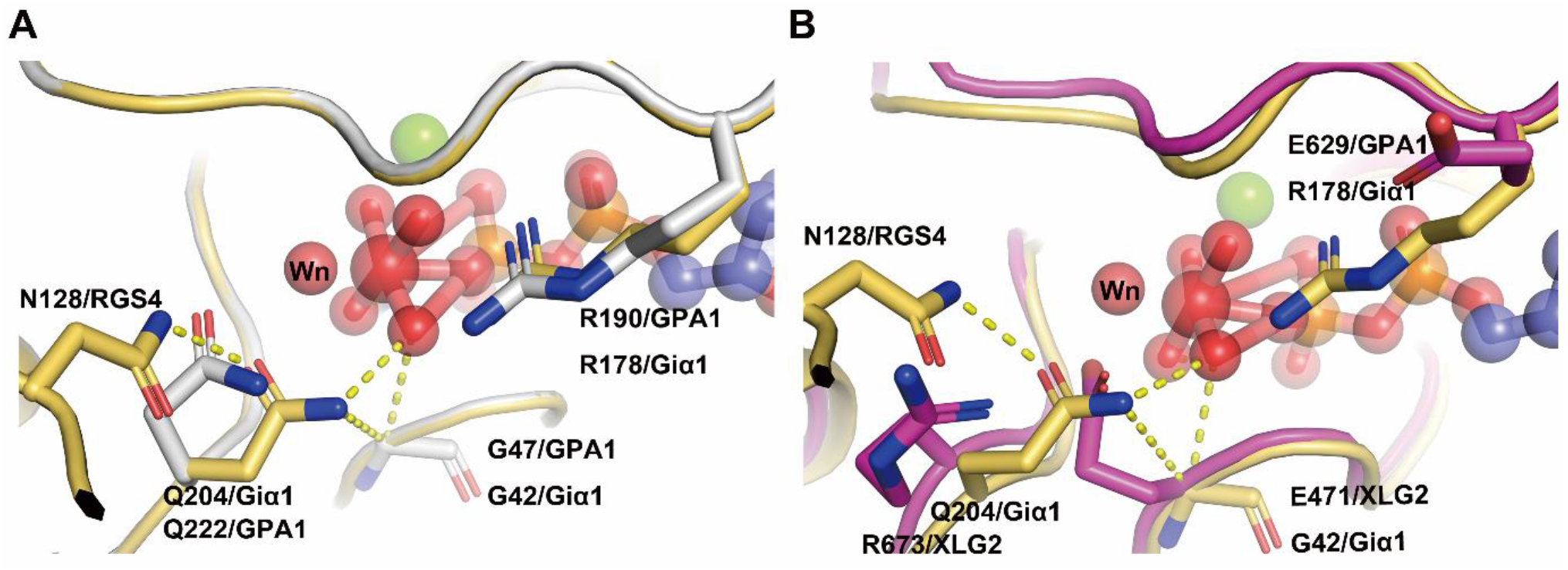
Interactions between the catalysis center of the Ras domain and critical residues of RGS proteins. **(A)** The critical contact residues between Gαi1 and RGS4 (PDB 1AGR) are shown in light orange. Arg 178 (Rcat) is within hydrogen bonding distance of the leaving group β-γ bridge oxygen and Q204 (Qcat) is a hydrogen bond donor to a fluorine (or O-) Al substituent and accepts a hydrogen bond from the presumptive water nucleophile (Wn). The hydrogen bond network (yellow dashed lines) involving N128 (Asn thumb of RGS4), Qcat, G42 and the the γ phosphate (modeled by AIF_4_) orient Wn for nucleophilic attack and stabilize developing charge at the β-γ bridge leaving group oxygen. RGS4 residues Asn 128 constrain the conformation of Gαi1 Q204 (Qcat) to the pre-transition state conformation. AtGPA1 contains the same catalysis network (A) however the catalysis network was disrupted in XLG2 with the loss of the Glncat and Arg finger and replaced by R673 and E629 respectively **(B)**. Grey: AtGPA1, Magenta: XLG2, Light orange: Giα1. The substrate GDP and AIF_4_ are shown as sticks and spheres. Main catalysis residues between Giα1, AtGPA1, XLG2 and RGS4 are highlighted as sticks. Wn: nucleophilic water.

The G3 box contains the signature sequence DXGG conserved throughout the heterotrimeric G-protein superfamily. Similar to the P loop, residues with the G3 motif interact with the γ-phosphate of GTP but also orients the Mg^2+^ ion that is critical for coordination of the guanine nucleotide. In AtGPA1 and Giα1, the G3 box is invariant “DVGG” (Fig. 2E), however in the XLG2 Gα domain, the residues are replaced by “RLNP” (Fig. 2F). The conserved Asp residue of canonical Gα subunits provides the water-molecule-mediated coordination of Mg^2+^ and therefore, the substitution of Asp for this critical Arg disrupts the ability to bind Mg^2+^ (6, 7, 27). Moreover, the main chain amide of the signature Gly residue is essential for nucleotide binding through hydrogen bonding to the γ-phosphate oxygen of the GTP (27). The main chain amide of this Gly is hydrogen bonded to the γ-phosphate and mutation of the two Gly residues in the G3 box confer dominant negative phenotypes (7, 27, 28). Gilman’s group showed that GDP-bound Gαs G226A mutant (the second Gly in the G3 DVGG motif) has a higher affinity for Gβγ than the wild-type subunit and is incapable of undergoing a GTP-induced conformational change (29). Taken together, stark differences in the three dominant residues in the G3 motif in the XLG2 protein nucleotide binding pocket which is conserved among all XLG proteins (Fig. 1) suggest that XLGs exist in the empty nucleotide state.

The G5 motif consensus is S/C/T-A-K/L/T. In Giα1, the G5 motif sequence is C-A-T (A = residue 326) and in AtGPA1, it is T-A-L (A = residue 355, Fig. 2G and H). In either form, the main chain of A326 in Giα1 is essential for the binding of GTP/GDP specifically forming a hydrogen bond with the oxygen of the guanidine nucleotide and S substitution at this site weakens the affinity for GTPγS through steric crowding (30). Also, the equivalent A366S mutation in the G5 motif of Gαs decreases Gαs’s affinity for GDP and GTPγS by steric crowding and shifting Gα towards the empty nucleotide pocket state (30, 31). However, in XLG2, the equivalent residues are C-Q-V (Q = residue 818, Fig. 2H). Thus, this substitution of A326 with Q818 in XLG2-1 is predicted to create a steric clash for nucleotide binding providing further inference that XLG is nucleotide-free.

### XLG proteins lack key residues to catalyze GTP hydrolysis

Two amino acids, one from the Gα subunit (the conserved catalytic glutamine residue in Switch II region which is named “Qcat”) and one from the RGS protein (the so-called “Asn thumb”), together with nucleophilic water and a Mg^2+^ in the catalytic center are essential elements for the catalytic reaction (6) (Fig. 3). In Giα1, the Qcat in Switch II is Q204 is essential for catalytic activity in the Gα subunit. A conserved Arg residue in Switch I region designated “Rcat” here, is also a major determinant of the catalytic activity. A water molecule designated “Wn” occupies the position for the nucleophile engaged in an in-line attack on the phosphate. The Asn thumb (N128 in RGS4) in the RGS domain reorients the Qcat allowing the carboximido moiety to form hydrogen bonds with AlF_4_ mimicking a γ phosphate oxygen atom and Wn. Rcat forms electrostatic interactions with the β phosphate oxygen and with one of the fluoride substituents of AlF_4_ (Fig.3). Mutations in these residues of Switch I and Switch II are known to drastically alter GTPase activity (6).

All XLG proteins lack both essential Rcat and Qcat for the catalysis (Fig. 3B). In XLG proteins, the Rcat residue in Switch I is E, creating a charge reversal that disrupts electrostatic interactions with the β and γ phosphates of the guanine nucleotide. The equivalent mutation in Gαi1 exist as a stable protein in a nucleotide-free state and lacks the capacity to form the active conformation (19). In all XLG proteins, the Qcat residue of Switch II is R/K which is unable to coordinate with either the Asn thumb of the RGS protein or the nucleophilic water to hydrolyze GTP. Both Q204R and R178C mutations abrogate nucleotide hydrolysis (19). The structural characteristic of the XLG proteins catalysis center suggests that XLGs lack the ability both to coordinate with RGS to hydrolyze GTP and the intrinsic GTPase activity of Gγ subunits.

### XLG2 has a much lower binding affinity towards nucleotide than canonical G subunits yet interacts with similar affinities towards Gβγ and AtRGS1

Assessments of nucleotide binding to XLG proteins to date lack quantitation for binding constants (32). Similarly, XLG protein interaction with AtRGS1 and Gβγ have been indirect measurements (11, 16, 17). To correct this deficit, we used microscale thermophoresis (MST) to measure the binding affinity of XLG2 and AtGPA1 with guanine nucleotides (GDP and GTPγS) and with binding partners AtRGS1 and Gβγ. The advantages of this new technique are the capability of obtaining accurate affinities in the low affinity (μM-mM Kd) range with small amounts of protein. Note that, unlike MST, traditional radioisotope binding assays are not accurate for low affinity interactions. Raw data with the quality control parameters provided are in Fig 4 and S3 and are summarized in Table 1. The observed Kd of AtGPA1 binding GTPγS was ~ 21 nM. This is within the range of Kds reported for animal G subunits (10-100 nM, (33)). XLG2 bound GTPγS with a Kd of ~ 2 μM which is 100 times lower affinity than GTPγS binds AtGPA1 when tested under the same conditions and nearly 1000 times lower when measured using radioactive ligand (34). Moreover, the affinity of XLG2 to GDP is ~100 fold lower (177 μM) than for GTPγS. Quantitative analyses clearly show that XLG2 is severely impaired in guanine nucleotide binding (Table 1).

**Table 1.** Summary of the binding affinity (Kd) among AtGPA1 and XLG2 with nucleotide, RGS1-C domain and Gβγ. Top row in the inset indicates the tested interactors. The values were determined from the binding isotherms shown in Figure 4. The values are averages and StdDev for all the experimental replicates. Each experiment was replicated at least once.

| | GTPγS | GDP | RGS1-C domain<br>Gα in apo state | RGS1-C domain<br>Gα in transition<br>state | G$\beta\gamma$<br>Gα in GDP state |
| --- | --- | --- | --- | --- | --- |
| AtGPA1 | 21 ± 18 nM | 28± 12 μM | 125 ± 81 nM | 67 ± 18 nM | 2 ± 1.22 μM |
| XLG2 | 2350 ± 460 nM | 177 ± 33 μM | 198 ± 45 nM | NA | 0.7 ± 0.09 μM |

**Figure 4.**
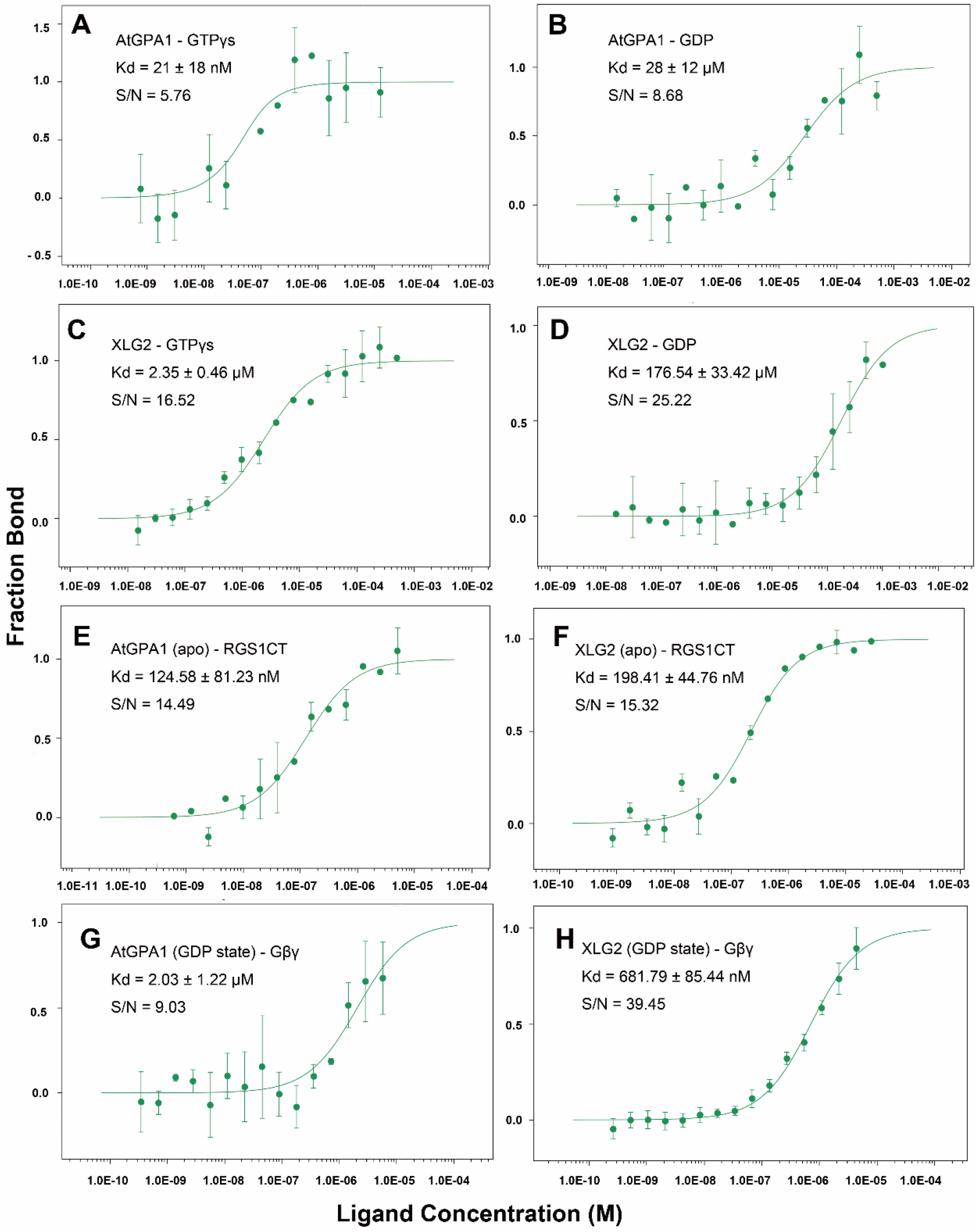
Binding isotherms for nucleotide, RGS1-C domain and Gβγ to AtGPA1 and XLG2. Microscale Thermophoresis was used. **(A)** Binding isotherm and Kd value of AtGPA1 binding GTPγS and **(B)** GDP. **(C)** Binding isotherm and Kd value of XLG2 binding GTPγS and **(D)** GDP. **(E)** Binding isotherm and Kd value of RGS1 C terminal domain to AtGPA1 apo state and **(F)** XLG2 apo state. **(G)** Gβγ binding to AtGPA1 and **(H)** XLG2 Gα domain. S/N: signal to noise ratio. Each experiment was repeated at least once. Binding curve and Kd were fitted as described in Methods. Error bars represent StdDEV. Each experiment was repeated at least once.

With this poor affinity toward guanine nucleotides, the concentration of GTP in plant cells would need to be 100 times greater than in animal cells for XLG2 to be GTP bound, however, for several reasons, this explanation of a mechanism to compensate the weak GTP affinity by XLG proteins is not reasonable. First, such a saturating concentration of GTP would eliminate the switch-like behavior of the canonical plant Gα subunit. Second, protein translation uses the same machinery in both plant and animal cells and the GTP hydrolyzed for its proof reading and is sensitive to its cytoplasmic concentration. Similarly, plant and animal microtubules requires GTP binding and hydrolysis. Both translation and cytoskeletal dynamics would cease at this high concentration of GTP. Third, nucleotide synthesis uses product inhibition to control the levels accordingly. A 100-fold higher concentration of GTP would be incompatible with enzymes involved in nucleotide synthesis. Fourth, the highest known concentration of GTP in plant cells is equivalent to only one Kd for GTP binding to XLG2 (35–37). As such, the concentration of GTP in plant cells, especially non-dividing cells, may be rate-limiting for full occupancy of XLG2 by GTP. For these reasons, we conclude that XLG2 is not likely bound by GTP *in vivo*.

Assmann’s group reported the unusual finding that the three XLG proteins bind and hydrolyze GTP using Ca^2+^ instead of Mg^2+^ as a coordinating factor (32). To test this, we performed MST experiments to measure the binding affinity of XLG2 with nucleotide in the presence of Ca^2+^. The results showed lower binding affinity towards GTPγS (~186 μM) with Ca^2+^ vs. Mg^2+^ (Fig S3). This indicates that Ca^2+^ may not act as the cofactor for XLGs binding GTPγS. Ca^2+^ induced relatively higher binding affinity for GDP (~28 μM), albeit still poor, compared to Mg^2+^ as a cofactor (Fig S3).

Interestingly, despite XLG2 having much lower binding affinity towards GTPγS and GDP compared with AtGPA1, it had a similar binding affinity to the C-terminal RGS domain of AtRGS1 and to the Arabidopsis Gβγ dimer (AGB1/AGG1). AtGPA1 bound AtRGS1 with a Kd ~125 nM with in AtGPA1 in its apo state and ~ 67 nM in its transition state (Fig.S3). XLG2 has a similar Kd of ~198 nM towards AtRGS1 when in its apo state (Table 1 and Fig. 4). The Kd for a transition state XLG2 was not determined because this state is not relevant due to its nucleotide independence. Moreover, XLG2 showed a ~0.7 μM binding affinity towards Gβγ similar to that of AtGPA1 which is ~2 μM (Table 1 and Fig. 4). This suggests that XLG2 exists as a nucleotide-independent inhibitor of G signaling through its ability to sequester Gβγ directly or indirectly by binding to AtRGS1 thus enabling freed AtGPA1 to sequester Gβγ.

### A mechanistic explanation: Relative instability of XLG2 confers the reduced nucleotide interaction

We applied several computational modeling and simulation approaches to understand the underlying molecular mechanisms differentiating AtGPA1 and XLG2 proteins. We sought to provide structural and molecular dynamics rationales for the experimentally observed differences in nucleotide binding preferences by the two proteins. To this point, we performed microseconds of molecular dynamics (MD) simulations of four molecular complexes, involving GDP and GTP nucleotides, each in complex with both AtGPA1 and the homology-modeled XLG2-1 Gα domain, followed by comparative analyses of the respective MD trajectories. The main finding of our simulations is that the molecular dynamic behaviors of XLG2-1 differs from that of AtGPA1. We observed that overall XLG2-1 was more mobile in comparison with AtGPA1, which generally retained its original crystallographic structure over the course of simulations. Furthermore, to distinguish the two proteins with respect to their nucleotide binding capabilities, we focused on analyzing the behavior of the ligand binding site both in the context of the intra-protein and ligand-protein interactions in order to more clearly understand the key factors contributing to the experimental findings of the lower nucleotide binding affinity in XLG2.

In preparation for MD simulations, the structure of XLG2 obtained by homology modeling, was subjected to molecular mechanics minimization following several protocols as described in the Methods section in order to avoid unnatural clashes between atoms resulting from homology modeling. To understand the overall dynamics of the proteins, we analyzed RMS fluctuations per residue and calculated RMSD using all C-alpha atoms of the proteins (**Fig. S4, S5**), which showed that the general fold of AtGPA1 was more stable and the amino-acid residues displayed lower mobility compared to XLG2. We then sought to understand the dynamics of the nucleotide binding site and explored the key differences in the interactions formed within the binding site. First, we visualized the binding sites of the two proteins to explore the main differences in terms of the amino-acid residue composition (**Fig. 5A**). The following differences in the similarly-positioned, binding-site residues were determined between AtGPA1 and XLG2: E48 to K472, D162 to R601, R190 to E629, F253 to E705, R260 to K714, K288 to K742 in guanine and ribose binding sites, and K51 to A475, S52 to T476, T193 to S632, D218 to R669, Q222 to R673 in Mg^2+^ and phosphates binding sites (**Fig. 1, 5**).

**Figure 5.**
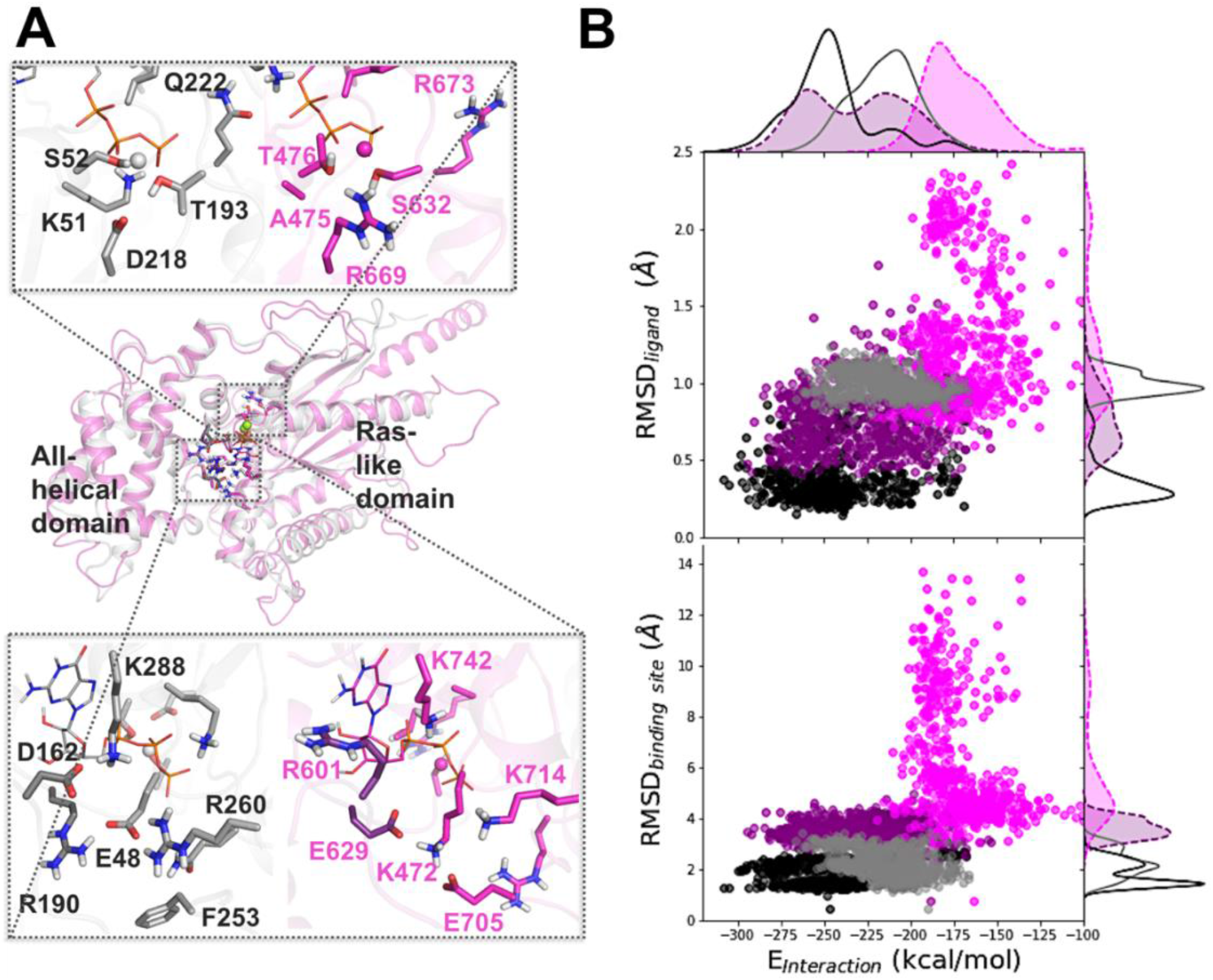
Difference in dynamics between the nucleotide-bound AtGPA1 and XLG2: Insights from MD simulations. Homology modeling and MD simulations reveal the main differences in amino-acid residue composition and the nucleotide binding site dynamics of AtGPA1 (grey) and XLG2 (magenta) **(A)** Aligned minimized AtGPA1 crystal structure (PDB ID 2xtz) and the homology modeled XLG2. The zoomed in plots separately display phosphate and Mg^2+^ binding site (top) and guanine and ribose binding site (bottom) on the example of GTP-bound complexes, highlighting the most prominent differences in the amino-acid residues. **(B)** The relationship between the nucleotide-protein interaction energies (designated as E_interaction_ on the scatter plots, and calculated as the sum of the Coulomb and LJ terms) and mobility of the nucleotide (RMSD*_ligand_*) and binding site (RMSD*_binding site_*) display substantial separation among AtGPA1-GTP (black points and solid line), XLG2-GTP (purple points and dashed line), AtGPA1-GDP (grey points and solid line), and XLG2-GDP (pink points and dashed line) complexes (Fig. S6-8).

To understand the differences in the binding site dynamics, we first calculated RMSD of the heavy atoms of residues located in the binding sites (**Fig. 5B and Fig. S6**), which we defined as the protein residues within 4 Å from GTP (see Methods). We observed that AtGPA1 and XLG2 nucleotide binding sites differed in conformational dynamics and had distinctly different configurations as elaborated in the following paragraph. Next, we aimed to understand the impact of the difference in dynamics of the binding site residues on the nucleotide mobility and nucleotide binding preferences (**Fig. 5B & Figs. S7, S8**). Through exploring the relationship between the nucleotide-protein interaction energy (calculated as the sum of intermolecular Coulomb and LJ terms of the molecular mechanics energy of the nucleotide-protein complexes) and mobilities of the binding site and ligand, we observed substantial differences across the four complexes formed when AtGPA1 and XLG2 bound to both GDP and GTP. The molecular systems occupied distinct regions on each of these two landscapes. Importantly the ranking order of the means of two parameters, (i) the nucleotide mobility in the pocket as characterized by the RMSD of the nucleotide (from smallest to largest), and subsequently (ii) the nucleotide-protein interaction energies characterized as the sum of all LJ and Coulomb terms of the nucleotide-protein interactions (from more negative to less negative), agreed with the ranking order in terms of our experimental binding affinities (Kd +/- StDev) as follows: **1^st^)** GPA1-GTP (0.021 +/- 0.018 μM), **2^nd^)** XLG2-GTP (2.4 +/- 0.5 μM), **3^rd^)** GPA1-GDP (28 +/-12 μM), **4^th^)** XLG2-GDP (177 +/-33 μM) (**Table 1**).

This result added confidence to our structural and simulations-derived interpretations of the molecular complex formations. We would like to emphasize, however, that such calculations of the intermolecular interaction energies are merely estimates of the relative strengths of ligand-protein interactions in the bound state, which by no means is equivalent to the assessment of the change in Gibbs free energy of binding (38–41).

Cluster analysis of the generated MD trajectories (42–44) (see Methods for details) revealed metastable states with distinct configurations of the binding sites linked to the experimental nucleotide binding preferences (**Fig. 6**). For the most populated metastable state of each molecular complex, we explored the specific intra-protein chemical interactions which directly impact the dynamics of the active sites, and their impact on the nucleotide-binding interactions. The results show that nucleotide-bound AtGPA1 achieves stable dominant (i.e., frequently visited) conformations, whereas XLG2 complexes tend to transition between conformationally-diverse states with lower probabilities. Such low frequency populations of the top clusters are associated with a more dynamic binding pocket in XLG2. Interestingly, both apo-proteins assume multiple states with low probabilities across the top five clusters (**Fig. S6A, S9, Supplemental Movies 1-6**).

**Figure 6.**
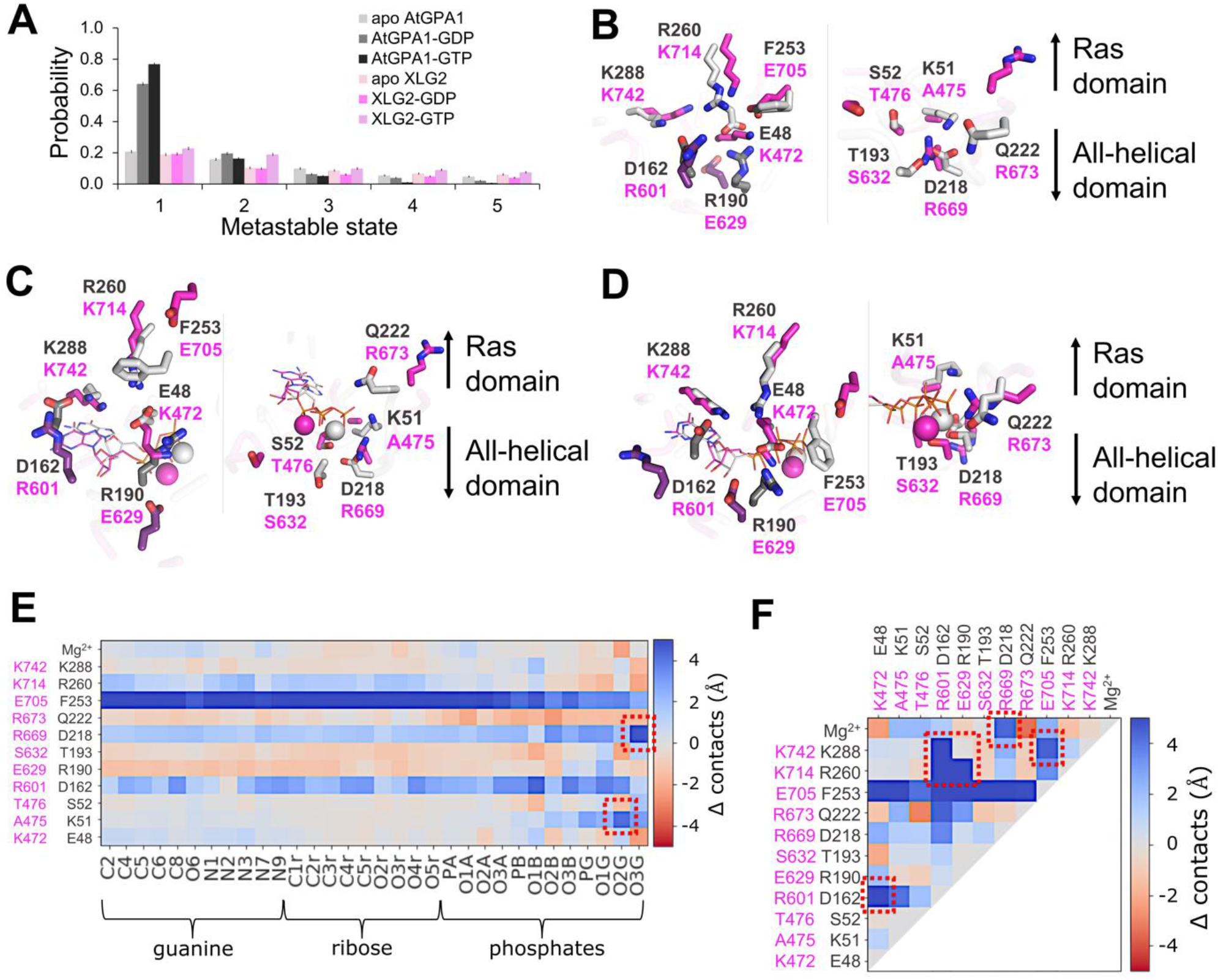
Apo and nucleotide-bound proteins obtain distinct configurations defining the nucleotide binding preferences of AtGPA1 and XLG2. **(A)** The top five most populated metastable states of the nucleotide binding site obtained in cluster analysis of MD trajectories indicate that nucleotide-bound AtGPA1 obtains a stable frequently visited conformational state, whereas XLG2 complexes tend to transition between conformationally diverse states with lower probabilities. Interestingly, both apo proteins obtain multiple states with equivalently low probabilities (Fig. S6,S9). Aligned centroids of the largest metastable states are presented in panels **B-D. (B)** The most populous apo states show stable E48-R190-R260 and D162-K288 salt bridge networks in the guanine binding site (left image) of AtGPA1 (grey), and a more destabilized salt bridge network between similarly positioned residues in XLG2 (magenta) primarily contributed by R601-K742 electrostatic repulsion. K51-D218 salt bridge in the phosphate and Mg^2+^ binding sites (right image) enables a more structures AtGPA1, while the neutral A475 and a repulsion between R669 and R673 cause a more disintegrated XLG2. **(C)** GDP- and **(D)** GTP-bound complexes retain the strong salt bridge network in AtGPA1 and less stable electrostatic interactions in XLG2. K51 reorients and forms an additional bond with phosphates in AtGPA1, which is prevented by the equivalently positioned neutral A475 in XLG2. K472 breaks its bonds with E629 and rearranges to interact with closer located phosphates. The absence of *γ*-phosphate in GDP makes the nucleotide more mobile, losing the frequency of its contacts. R673 in GTP-bound XLG2, however, forms a relatively stable bond with the *γ*-phosphate seemingly increasing the nucleotide binding affinity. The residues shown in darker shades in panels B-D (D162 and R190 of GPA1; R601 and E629 of XLG2) make the key intra-protein interactions defining the binding site shape. The differences in the frequency of the aforementioned interactions, on the example of GTP-bound complexes, are clearly seen through heatmaps of Δ contacts (minimum distances) **(E)** between the non-hydrogen atoms of the nucleotide and binding site residues, and **(F)** within the binding site residues. 100 states of the most populated clusters were used to generate the heatmaps. The red squares highlight the most prominent changes in the interactions stipulating the importance of the residues to the stability of the active sites and to the interactions with the nucleotide.

The most populous states of the apo-protein show a stable salt bridge network in AtGPA1, but no network and fewer coherent salt bridges are found in XLG2-1. In the guanine binding site of AtGPA1, salt bridges formed between E48 of the P-loop, R190 of Switch I, and R260 of Switch III, as well as between D162 and K288, while a more destabilized salt bridge network appeared between similarly positioned residues in XLG2-1, namely: E629 of Switch I, K472 of P-loop, K714 of Switch III and E705 (**Fig. 6B, S9-12**). In XLG2-1, a positive charge at K714 (position equivalent to R260 in AtGPA1) is not capable of forming a salt bridge with K472 (position equivalent to E48 in AtGPA1). In the phosphate and Mg^2+^ binding sites, the K51-D218 salt bridge enables a more structured apo-AtGPA1, while for XLG2, the neutral sidechain of A475 (position equivalent to K51 in AtGPA1) together with a repulsion between R669 and R673 precludes a stabilizing salt bridge. In general, AtGPA1 salt bridges formed by the two loop residues D162 and R190 (and equivalently placed E629 in XLG2) drawing the two domains closer together to subsequently increase the number of interactions between the all-helical domain and the nucleotide.

The nucleotide-bound complexes retain the aforementioned strong salt bridge network in AtGPA1 and weak electrostatic interactions in XLG2 (**Fig. 6C,D, Fig. S9-12**). In the bound state, the K51 sidechain of AtGPA1 was re-arranged to form an additional bond with phosphates, which is an additional salt bridge lacking by the neutral A475 in XLG2-1. In the GTP-bound XLG2-1 model, E705 lost its K472 interaction to *γ*-phosphate which also attracted R673 for further stabilization. Moreover, E48 in AtGPA1 avoided the negatively-charged phosphates which further promoted electrostatic interactions with both R190 and R260 to stabilize this cavity. In the GDP-occupied state, due to the lack of the *γ*-phosphate, GDP is more mobile and loses a number of its contacts with the active site residues in both Gα proteins, but more so in XLG2-1 due its structurally-unstable binding site.

The heatmaps of Δ contacts from experiments determining the difference in the minimum distances between the two Gα proteins and **(i)** the atoms of the nucleotide and binding site residues, and **(ii)** intra-protein interactions within the binding site residues, clearly show the contrast between the interaction frequencies within these two proteins. As highlighted in Fig 6E and F, the most prominent changes in the interactions important to the stability of the active sites and to the interactions with the nucleotide, in addition to the previously indicated interactions, reside with F253 in AtGPA1 which was overall closer to both the nucleotide and the binding site residues compared with similarly positioned E705 in XLG2-1. This aromatic residue makes frequent pi-cation interactions with R190 in both apo- and ligand-bound protein and occasional pi-cation interactions with the Mg^2+^ in the ligand-bound state.

Our analyses shows that the D162-K288 salt bridge (**Fig. 7A**) is one of the key interactions maintaining the shape of the apo-AtGPA1 nucleotide-binding site. In contrast in XLG2-1, two positively charged residues, R601 and K742, situated in positions equivalent to D162 and K288 of AtGPA1 caused electrostatic repulsion, pushing away the all-helical domain of the protein from the Ras-like domain, resulting in an increased mobility and a less structured nucleotide binding site in XLG2-1. The R601-K742 distance in XLG2-1 was highly correlated with the fluctuations of the binding site in XLG2-GDP and to a lesser extent in XLG2-GTP (**Fig. 7B and C**). The reason for the former is the lack of an extra anchor in terms of *γ*-phosphate in GDP to enable the electrostatic repulsion between R601 and K742 to be the main contributor to the instability of XLG2-GDP binding pocket. This agrees with the experimentally observed poor binding affinity of GDP to XLG2. Although this correlation still exists in XLG2-GTP, it is less pronounced due to the presence of the extra phosphate in GTP. In XLG2, K714 makes ionic bonds with phosphates, however, this does not seem to be sufficient to retain the binding site integrity distorted by the aforementioned repulsion.

**Figure 7.**
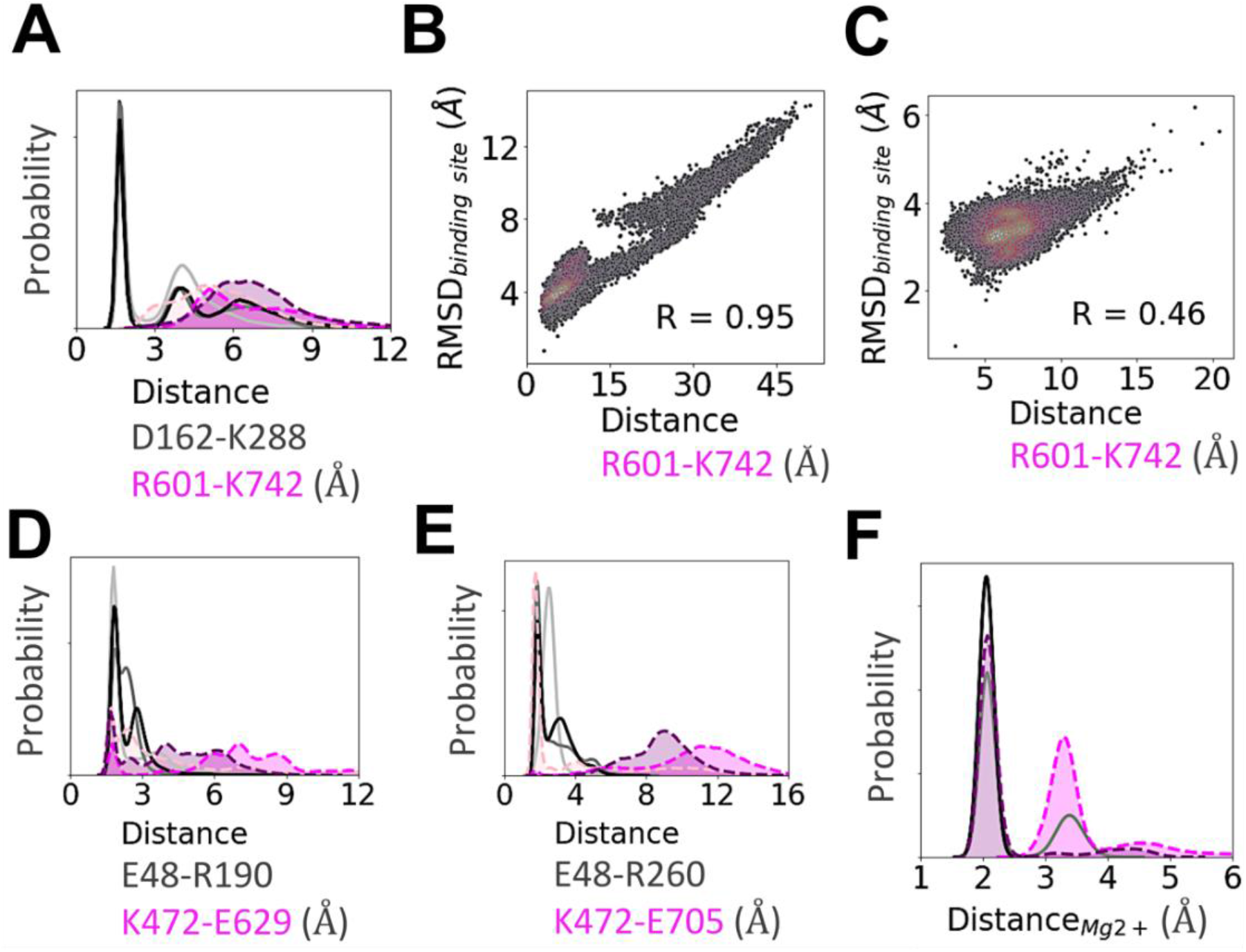
Key intra-protein distances responsible for the experimental nucleotide binding affinities as determined from MD simulations. Distribution of the minimum distances between the residues emphasized in Fig. **6** and text. In general, a stronger salt bridge network in AtGPA1 maintains the shape of its nucleotide-bound binding site (see text for more details). **(A)** Distance between D162 and K288 and the equivalently placed R601 and K742 in XLG2. **(B)** The repulsion between R601 and K742 is highly correlated (R=0.95) with the binding site RMSD in XLG2-GDP and **(C)** to a lesser extent in XLG2-GTP (R=0.46). The 2D correlation plots were constructed using kernel-density estimation with Gaussian kernels. **(D)** Distance between E48 and R190 in AtGPA1 and similarly positioned K472 and E629. **(E)** Distance between E48 and R260 in AtGPA1 and K472 and E705 (aligned with F253 of AtGPA1) in XLG2. The distributions show that both of the salt bridges (panels E and D) are dominant in the apo proteins, and are less persistent in nucleotide-bound XLG2. **(F)** The distributions of the minimum distance between Mg^2+^ and *γ*-phosphate binding site residues (D218, S52, T193, and D218 in GPA1; T476, S632, R669 in XLG2) and Mg^2+^ counterion. For clarity in panels A, D, E, and F the values of probabilities on the y-axis are hidden. Apo AtGPA1 is plotted with solid light grey lines, AtGPA1-GDP— solid grey, AtGPA1-GTP—solid black, apo XLG2—dotted pink, XLG2-GDP—dotted magenta, XLG2-GTP— dotted purple; the probability densities for XLG2 are shaded for contrast.

Another distinction between the two proteins is in the Mg^2+^ binding site (**Fig. 7F & Fig. S13**) discussed above. In AtGPA1, D218 formed an H-bond with S52 (which interacts with Mg^2+^) and a Coulomb interaction with Mg^2+^, whereas the ‘bulkier’ and positively charged sidechain of R669 in XLG2 (position equivalent to D218 in AtGPA1) did not form stable interactions with either T476 or S632 (which interact with Mg^2+^) and caused an electrostatic repulsion with Mg^2+^. The distribution of the minimum distance between the Mg^2+^ binding site residues (S52, T193, D218, and Q222 in AtGPA1; T476, S632, R669, and R673 in XLG2) and the Mg^2+^ counterion clearly explain this effect.

To interpret the observed equivalent AtRGS1 binding capability of the two Gα proteins (**Fig. 3, 4, Table 1**), we estimated the structural stability of the specific regions that are involved in AtRGS1 binding (**Fig. S14**). We showed that the three equivalently placed AtRGS1 binding site residues of apo-AtGPA1 and apo-XLG2-1 similarly maintained their structural integrity over the course of our simulations. Such conformationally-preserved regions in the apo-proteins position them to bind AtRGS1 when it is tethered close to either protein.

## Conclusion

XLG2 binds GTP *in vitro* poorly such that at the estimated concentration of GTP in plant cells, XLG2 is not expected to be nucleotide bound. However, XLG2 binds regulatory partners, AtRGS1 and Gβγ. Therefore, XLG2 is a decoy that negatively regulates by sequestering the Gβγ dimer directly and also indirectly by promoting AtGPA1 interacting with Gβγ through freeing AtGPA1 from the AtRGS1::AtGPA1 complex. While this concept shares similarities for control of G signaling by dominant negative mutations of canonical G protein in animals (45), it is unique in that the negative control is provided *in trans* by a genetically-encoded, atypical G protein.

Taken together, our modeling data provide credible interpretations for the experimentally observed strengths of guanine nucleotide binding to AtGPA1 and XLG2. Several key intra-protein and nucleotide-protein interactions in AtGPA1 were shown to be attributed to the higher structural stability of the binding site of the protein and to more persistent contacts of the protein with the nucleotide and magnesium. We show mechanistically that among the chief intra-protein interactions preserving the stability of the binding site in both apo- and nucleotide-bound-states of AtGPA1 include the following ionic bonds: D162-K288, R190-E48-R260, and K51-D218. Because XLG2 is important for disease resistance and development (17, 46, 47), engineering these equivalent residues may lead to improvements in crop performance.

## Acknowledgments

This work was supported by NIGMS (R01GM065989) and NSF (MCB-0718202) awarded to Alan. M. Jones. The deep computational analyses were supported by the resources of the UNC Longleaf cluster (https://its.unc.edu/research-computing/longleaf-cluster/)

