## Supplementary material for "A decoy heterotrimeric Gα protein has substantially reduced nucleotide binding but retains nucleotide-independent interactions with its cognate RGS protein and Gβγ dimer": materials and methods

### Protein expression and purification

AtGPA1 and XLG2 proteins were expressed and purified as described previously (47, 48). For protein expression, proteins were transformed into ArcticExpress RP cells (Agilent Technologies). In large scale culture, 0.5 L LB medium in 2.5 L flasks were incubated at 37 ºC, 225 rpm. At OD600 = 0.6 to 0.8, protein expression was induced using 0.5 mM IPTG at 12 ºC for 16 hours. All purifications were performed at 4 ºC.

### Purification of AtGPA1 and XLG2

For quantification of interaction between AtGPA1 and AtRGS1 and AGB1/AGG1, Twinstep-GPA1 was used. Twinstrep-AtGPA1-pDEST17 (His tag removed) was transformed into ArcticExpress RP cells (Agilent Technologies). Protein expression was induced by 0.5 mM IPTG at OD600 0.75 and cultured at 12 °C for 16 hours. Cell pellets were resuspended in extraction buffer (25 mM Tris-HCl pH 8.0, 150 mM NaCl, 2 mM $MgCl_2$, 20 µM GDP, 5 mM 2-mercaptoethanol, 1 mM PMSF, 0.25 mg/mL Lysozyme, 0.1% Thesit (Sigma, 88315), 1 X protease inhibitor cocktail, 10% glycerol) and mixed for 30 min at 4 °C. The suspension was sonicated (Sonic Dismembrator, Model 550, Fisher Scientific, power level 5, 0.50/0.50 off for 1 min, 2 cycles) to disrupt the cells. The lysate was centrifuged at 30,000 X g for 40 mins, and then the soluble fraction was collected and incubated with strep-tactin sepharose (50% suspension, cat no. 2-1201-010, IBA) for 30min at 4 °C. Then resin was washed with washing buffer (= extraction buffer above except with neither lysozyme nor thesit) and eluted with elution buffer (Washing buffer supplemented with 2.5 mM desthibiotin (Sigma-Aldrich)). The eluted proteins were run on size exclusion column (Superdex 200 10/300 GL, GE Healthcare) with running buffer (20 mM Tris-HCl, pH 7.5, 50 mM NaCl, 10 mM $MgCl_2$, 50 µM GDP, 1 mM DTT, and 10% Glycerol). Aliquoted protein samples were snap frozen by liquid nitrogen and store at -80 °C.

For quantification of interaction between AtGPA1 and guanine nucleotides, polyHis-tagged-AtGAP1 was used. For polyHis-tagged AtGPA1 production, the cells were resuspended in N1 buffer (25 mM Tris-HCl pH 7.6, 20 µM GDP, 100 mM NaCl, 5% glycerol, 10 mM imidazole, 10 mM $MgCl_2$, 12.5 mM 2- mercaptoethanol, 1 mM PMSF, 10 mM leupeptin, 0.25 mg/mL Lysozyme, 0.1% Thesit (Sigma, 88315)) and mixed for 30 min. After sonication (Sonic Dismembrator, Model 550, Fisher Scientific, power level 5, 0.50/0.50 off for 1 min, 2 cycles), the concentration of NaCl was raised to 300 mM and the lysate was mixed for another 30 min. The soluble fraction was separated by centrifugation at 30,000 × g for 45 min, then the supernatant was incubated with TALON Metal Affinity Resin (50 µL 50% slurry per 1 g cell pellet) for 1.5 h. The resin was washed with washing buffer (25 mM Tris-HCl pH 7.6, 300 mM NaCl, 5% glycerol, 10 mM imidazole, 10 mM $MgCl_2$, 10 mM leupeptin) and eluted with elution buffer (50 mM Tris-HCl pH 7.6, 300 mM NaCl, 5% glycerol, 300 mM imidazole, 10 mM $MgCl_2$). The eluate was dialyzed against dialysis buffer (20 mM Tris-HCl pH 7.6, 50 mM NaCl, 1 mM $MgCl_2$, 1 mM DTT) overnight.

The purification procedure for XLG2 was similar to His-tagged AtGPA1 purification except the buffer was pH 7.5 and the polyHis tag was cleaved by TEV protease to generate untagged XLG2.

His-and GST-tagged RGS+Ct (AtRGS1, residues from 284 to 459) were cloned into pDEST17 or pDEST15 destination vector as previously described (49, 50). The procedure for expression and purification was similar to His-tagged AtGPA1 purification except in the absence of GDP. The His-GST tag was cleaved by TEV protease. This recombinant TEV enzyme had a 100 percent cutting efficiency.

**Gβγ Protein Purification**

AGB1 and AGG1 were purified as described previously (51). For expression of Gβγ in Sf9 cells with the baculovirus expression system, N-terminal 6XHis tagged AGB1 and 6XHis tagged AGG1 were subcloned into the pFastBacDual vector (Invitrogen) at the *BamHI/PstI* and *XhoI/KpnI* sites, respectively. AGB1/AGG1 recombinant proteins were expressed by P2 baculoviral stock (titer ~ 2X $10^8$ pfu/ml) infection of sf9 cells at multiplicity of infection (MOI) of 1 for 48 hours. Purification steps were performed at 4°C. Pellets were resuspended in extraction buffer (25 mM Tris, pH 8, 200 mM NaCl, 20 mM imidazole, 1 mM 2-mercaptoethanol, 1 mM PMSF, 0.25 mg/mL Lysozyme, 0.1% thesit (Sigma, 88315), 1X protease inhibitor cocktail). The cells were disrupted by sonication (Sonic Dismembrator, Model 550, Fisher Scientific, power level 5, 0.50/0.50 off for 1 min, 2 cycles). Sonicated lysate was centrifugated at 50,000 X g for 30 mins to obtain the soluble fraction, and then the soluble fraction was incubated with Ni-NTA Agarose (Qiagen, Mat. No, 1018244) for 1 hour. The Ni-NTA Agarose was washed with washing buffer (same as extraction buffer with no lysozyme and thesit) and eluted with elution buffer (25 mM Tris, pH 8, 200 mM NaCl, 250 mM imidazole, 1 mM 2-mercaptoethanol, 1 mM PMSF, 1Xprotease inhibitor cocktail, 20% glycerol). The eluate was run on a size exclusion column (Superdex 200 10/300 GL, GE Healthcare) with running buffer (20 mM Tris-HCl, pH 8, 50 mM NaCl, 1 mM DTT, and 10% Glycerol). Aliquoted protein samples were snap frozen in liquid nitrogen and store at -80 °C. Purified proteins were examined by SDS-

PAGE gel with both Coomassie Blue staining and anti-His tag western blot. Aliquoted protein samples were snap frozen by liquid nitrogen and store at -80 ºC.

**Quantitative binding analyses by microscale thermophoresis (MST)**

Measurements for equilibrium binding were performed using 50 nM fluorescently labelled protein using Monolith NTTM Protein Labeling Kit RED-NHS (Nanotemper Technologies). The dye/protein ratio used was 10:1. Experiments were conducted at 25°C and in 'MST Buffer' [PBS pH 7.4, 0.05% Tween-20]. Protein sample was centrifuged for 30 min (21,000 × g, 4°C) before experiment. For thermophoresis measurements, ligand and labeled protein sample was mixed 1:1 with each of the ligand dilution series. After 10-min incubation at room temperature, each dilution was filled into Monolith NTTM MST Premium-coated capillaries (Nanotemper Technologies). A capillary scan was performed with 40% LED power. Binding curves were fitted to two sets of replicates. For binding experiments, Kd values were calculated via the MO. Affinity Analysis V2.3 software. This software performs a quality control based on the levels of starting fluorescence, and the distribution of the signal within the capillaries, an indicator of non-specific binding. A signal to noise ration above 5 provides high confidence of the data.

**XLG2 model building**

To compare the protein structures between XLGs and canonical Gα, we created high-quality models of the Gα homology domains of XLG1, XLG2 and XLG3 using MODELLER and the aligned sequences shown in Fig. 1. The human RGS4 and Giα1 transition state (Ligand: AlF4 and GDP) complex (PDB [1AGR]) was used as template to generate the models of XLG2. Five models (XLG2-1 to XLG2-5) were created using the *Automodel* script based on the template of human Giα1 (PDB [1AGR]). For evaluation and selection of the best model, we calculated the objective function (molpdf), Discrete Optimized Protein Energy (DOPE) score and GA341 assessment score between the model and the template (Figure S2). The first model XLG2-1 was selected given the relatively low value of the molpdf and overall DOPE assessment scores. In addition, DOPE scores were calculated per-residue and the template and the models were compared using Gnuplot (Figure S2).

**Molecular dynamics simulations**

The crystal structure of Arabidopsis AtGPA1 (PDB ID 2XTZ) (10) and the homology model structure of Arabidopsis XLG2 were used as the starting structures in the molecular dynamics (MD) simulations. Upon the alignment of the two proteins, the nucleotide from the 2XTZ structure of AtGPA1 protein was copied into the XLG2 protein. Histidine residues were protonated using the Interactive Optimizer in the H-bond assignment section of the Protein Preparation Wizard module available through Maestro (Maestro Schrödinger software (release 2018-4, Schrödinger, LLC: New York, NY)). The protein structures were then energy-minimized using OPLS3 force field (FF) and the Optimal scheme available in the Macromodel Minimization module of Maestro, which used PRCG method with 3-point line searcher since the number of unfixed atoms was more than 1,000. The molecular systems were then prepared to perform MD simulations in Gromacs 2018.2 simulation package using CHARMM36 protein forcefiled (52). End caps were added to both termini of each protein. The protein complex was minimized in vacuum using the steepest decent algorithm for 5,000 steps or until the maximum force of 1,000 kJ*mol-1*nm-1 was reached. The molecular systems were then solvated in TIP3P water (53), counterions were added for system neutrality, and NaCl was added by replacing water molecules in order to mimic 0.15 M physiological conditions. The total system sizes were ~80,000 atoms for GPA1 systems and ~130,000 atoms for XLG2 systems. Solvent energy-minimization was then performed, followed by a two-step equilibration, during which all heavy atoms of the system, excluding those of water and counterions, were restrained: 0.1 ns in NVT ensemble using the modified Berendsen thermostat (54) set at constant 310 K, and 1 ns in NPT ensemble at constant 1 atm and 310 K using the Parinello-Rahman pressure coupling (55). All simulations were conducted using the Leapfrog integrator in periodic boundary conditions. The 6-12 Lennard-Jones potential was used to describe the vdW interactions, and the nonbonded cutoff distance was set at 0.1 nm. The particle mesh Ewald algorithm (56) controlled the long-range electrostatic interactions. Bonds involving hydrogen atoms were constrained using the linear constraint solver algorithm (LINCS) (57). The production simulations were conducted in NPT ensemble with all atoms free to move. The volta GPU nodes on UNC Longleaf supercomputer cluster were used, and each simulation was performed on a combination of 1 GPU and 4 associated CPUs. Each of the six molecular systems were subjected to three independent 1,000 ns long MD runs totaling 18 μs cumulative simulation time. Gromacs's trajectory analysis tools, MDTraj (58, 59) along with *in house* bash and python scripts were used for data analysis, and matplotlib and seaborn were employed for plotting. Molecular visualization and generation of graphics were performed in MacPyMOL v1.8.6.2 (The PyMOL Molecular Graphics System, Version 1.8.6.2, Schrödinger, LLC). Cluster analysis was performed with gmx cluster available in Gromacs

(43) using the RMSD cutoff of 2.0 Å. The following binding site residues were used to calculate RMSD of the binding site and for clustering: GPA1: 47-53, 162-163, 187-193, 218-222, 253, 260, 287-288, 290-291, 354-356; XLG2: 471-477, 601-602, 606, 626-632, 669-673, 705, 714, 741-742, 744-745, 817-819. Additionally, for clustering of the nucleotide-bound complexes the ligand and magnesium were also included. For Fig. 6A the binomial standard deviations $\sigma$ of the state probabilities were calculated through

$$\sqrt{P(1-P)/N},$$

where *P* is the probability of the cluster and *N* is the total number of observations.
