## supplemental figures and legend for "A decoy heterotrimeric Gα protein has substantially reduced nucleotide binding but retains nucleotide-independent interactions with its cognate RGS protein and Gβγ dimer"

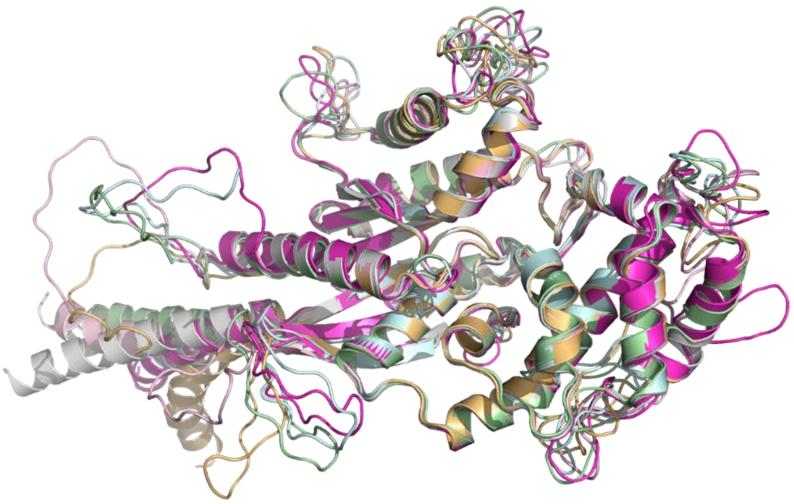

| Filename | molpdf | DOPE score | GA341 score |
|---|---|---|---|
| XLG2-1.pdb | 2721.09570 | -41152.59375 | 1.00000 |
| XLG2-2.pdb | 2605.27905 | -40820.97656 | 1.00000 |
| XLG2-3.pdb | 3669.00073 | -40187.55078 | 1.00000 |
| XLG2-4.pdb | 3337.97363 | -40364.93359 | 1.00000 |
| XLG2-5.pdb | 3192.82910 | -41316.17188 | 1.00000 |

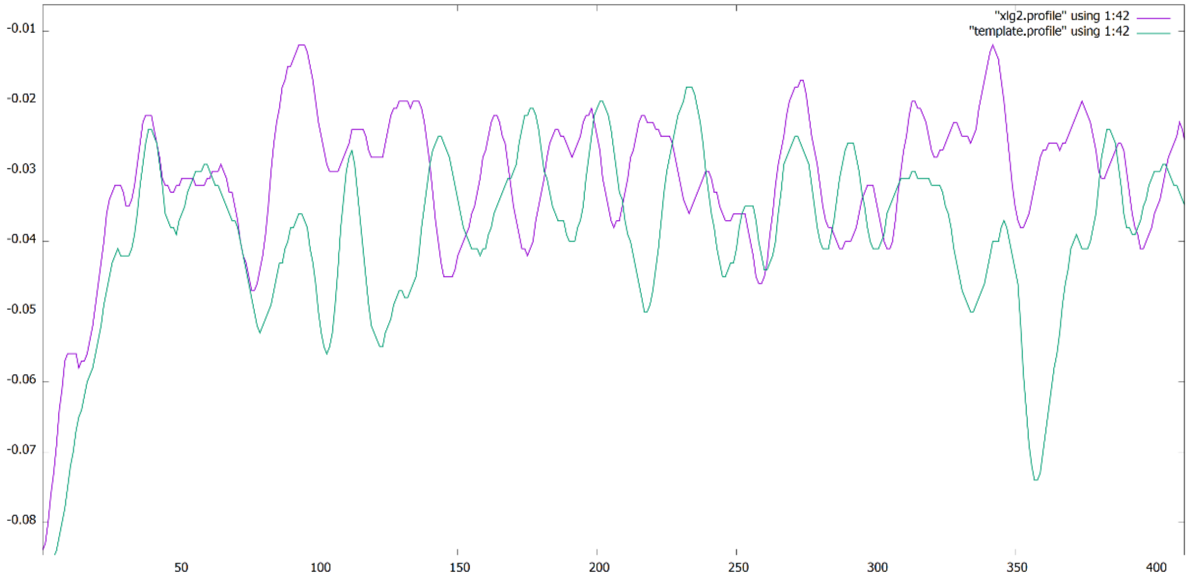


**Figure S1. Model evaluation and alignment results of the 5 models of XLG2 with Giα1 template (PDB[1AGR])** The results showed that the 5 models share overall similar fold with the template. Grey: Giα1 template, magentas: XLG2-1, Pink: XLG2-2, Pale-green: XLG2-3, Palecyan: XLG2-4, Light orange: XLG2-5The MODELLER objective function (molpdf), DOPE assessment scores (Discrete Optimized Protein Energy, which is a statistical potential used to assess homology models in protein structure prediction) and GA341 assessment score (range from 0.0 (worst) to 1.0 (native-like)) were calculated to evaluate the models. The first model XLG2-1 was selected given the lowest value of the molpdf and relatively low overall DOPE assessment scores. In addition, DOPE per-residue scores of the selected XLG2-1 model were calculated and were compared using Gnuplot.

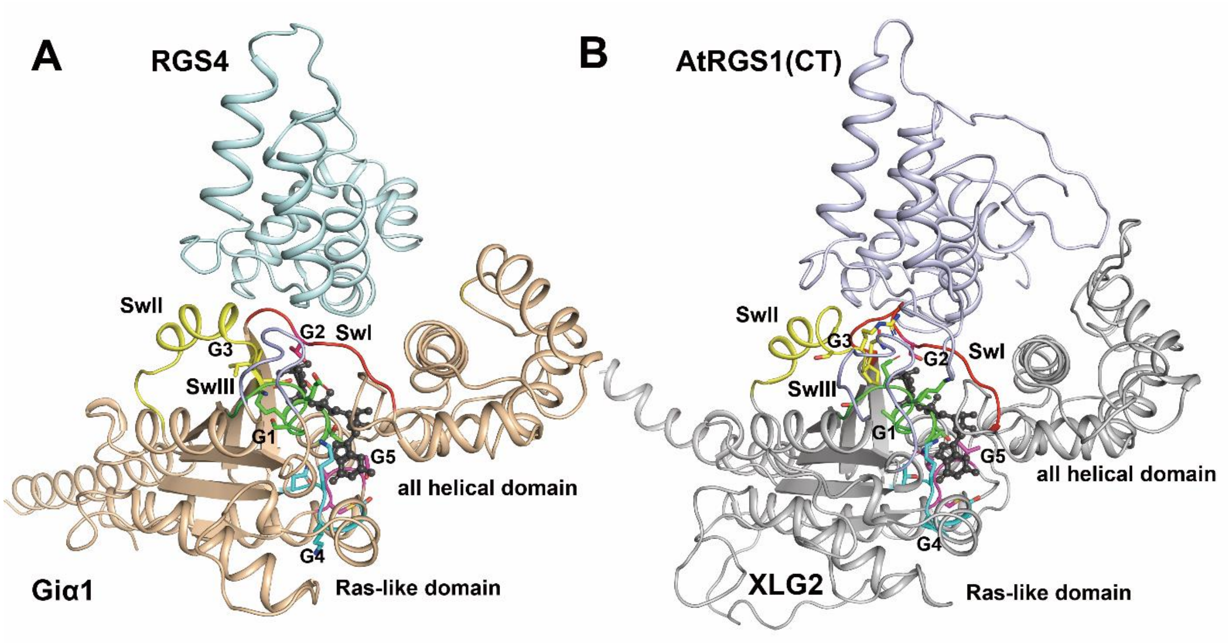


**Figure S2 Comparison of the model of XLG2 Gα homology domain in complex with RGS domain of AtRGS1 and the RGS4- Giα1 complex (PDB [1AGR])** (A) Crystal structure of the RGS4- Giα1 complex (PDB [1AGR]). RGS4 is colored in cyan and Giα1 colored in lightorange. (B) The model of the XLG2-1 in complex with RGS domain of AtRGS1. RGS1 is colored in light blue and XLG2 colored in grey. The substrate GDP and $AlF_4$ are shown as sticks and spheres and are colored in black. Switches I-III regions (SwI-III) are highlighted in red, yellow and light blue, respectively. The G1-G5 motifs are highlighted with different color and showed as sticks (G1 motif/P loop: green, G2 motif: red, G3 motif: yellow, G4 motif: cyan, G5 motif: magenta)

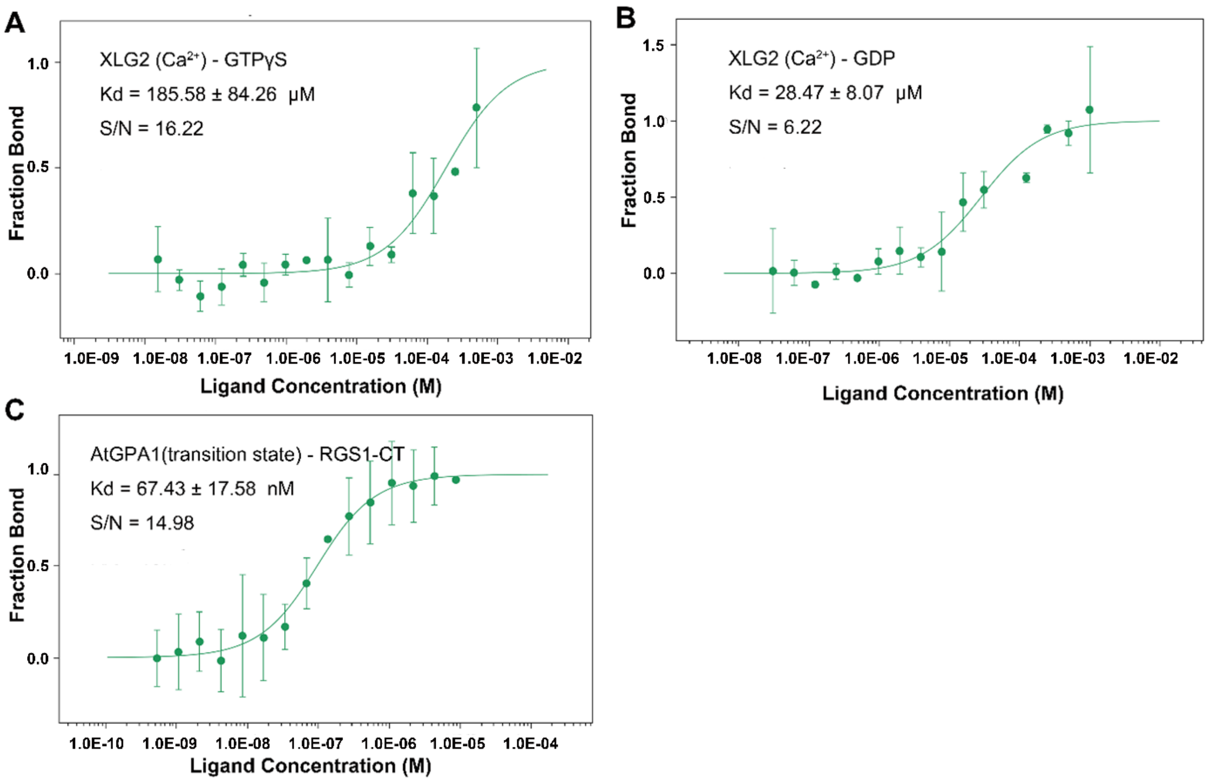


**Figure S3. MST binding results of XLG2 with GTPγs, GDP using Ca2+ as coeffector and AtGPA1 transition state with RGS1 C terminal domain** (A) Binding curve and Kd value of XLG2 binding GTPγs (B) Binding curve and Kd value of XLG2 binding GDP (C) Binding curve and Kd value of AtGPA1 transition state binding RGS1 C terminal domain (S/N: signal to noise ratio, RA: response amplitude. Each experiment was repeated at least once. Binding curve and Kd were fitted using MO. Affinity Analysis software.)

The following data were computed from the molecular dynamics simulations. Median values of the parameter (and median absolute deviation in the brackets) for each of the three independent simulation runs (seeds) of each simulated system (GPA1-apo, GPA1-GDP, GPA1-GTP, XLG2-apo, XLG2-GDP, XLG2-GTP) are indicated in the figure legends. The full data calculated from frames saved with 40 ps frequency is plotted using transparent lines, and a ‘clearer’ timeseries line is also plotted, which was obtained by using the Savitzky-Golay filter with window size 161 and polynomial order of 3 to remove high frequency noise from the data.

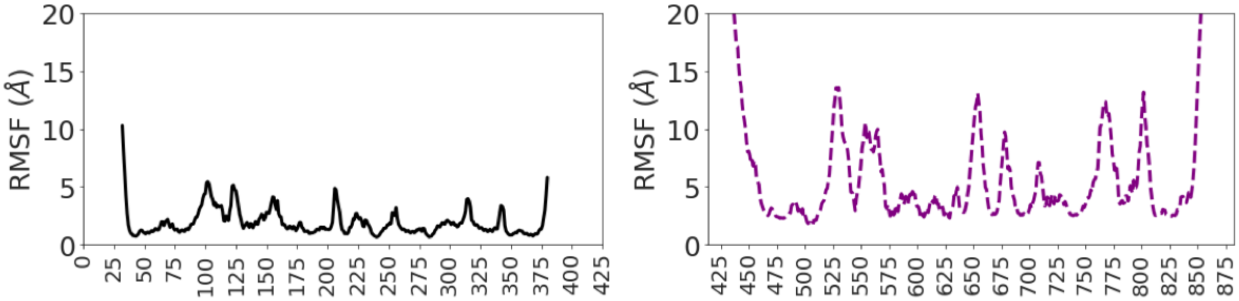


**Figure S4.** RMS fluctuations on the example of GTP-bound GPA1 (left) and XLG2 (Right) complexes. The reference structures were the minimized crystal structure for AtGPA1 and minimized homology model of XLG2. The plots indicate that while the overall trend of the fluctuating regions of the proteins are similar across both AtGPA1 and XLG2, XLG2 displays much higher fluctuations in comparison with AtGPA1.

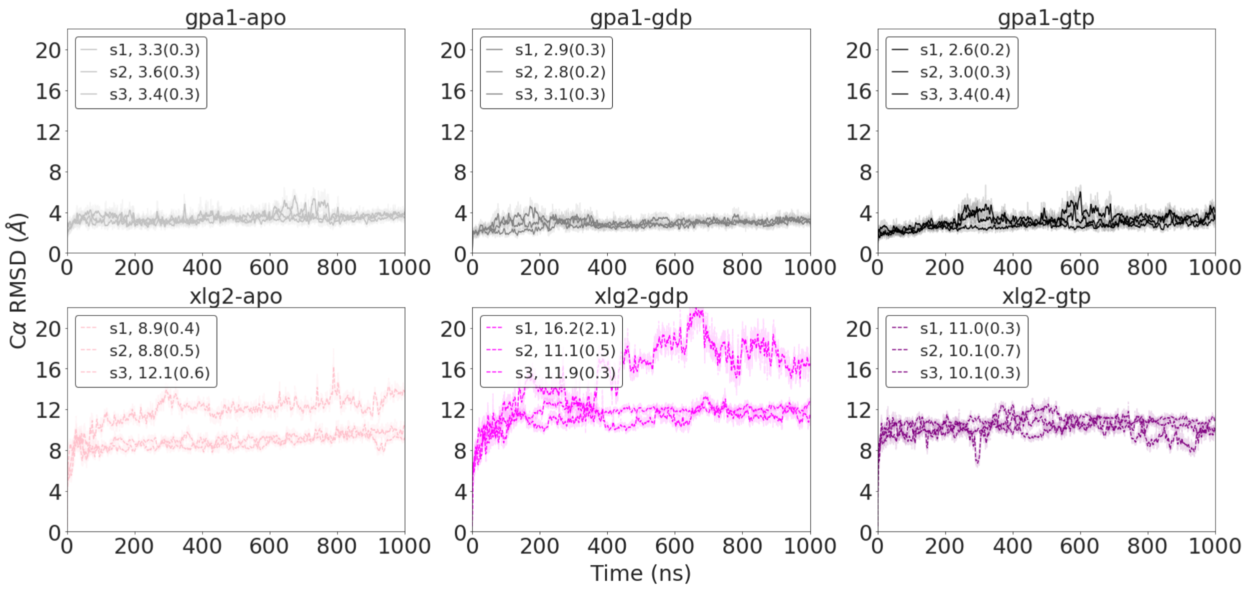


**Figure S5**. Timeseries of the full protein length C-alpha RMSD. The reference structure for GPA1 was the minimized crystal structure, while for XLG2 the minimized homology model. The data indicates that the overall structure of apo, GDP- and GTP-bound AtGPA1 remains stable throughout the simulations with low deviations. XLG2-GTP structure stabilizes in a relatively stable conformation, whereas apo and GDP-bound states possess higher mobility and obtain various configurational states (see the main manuscript for details).

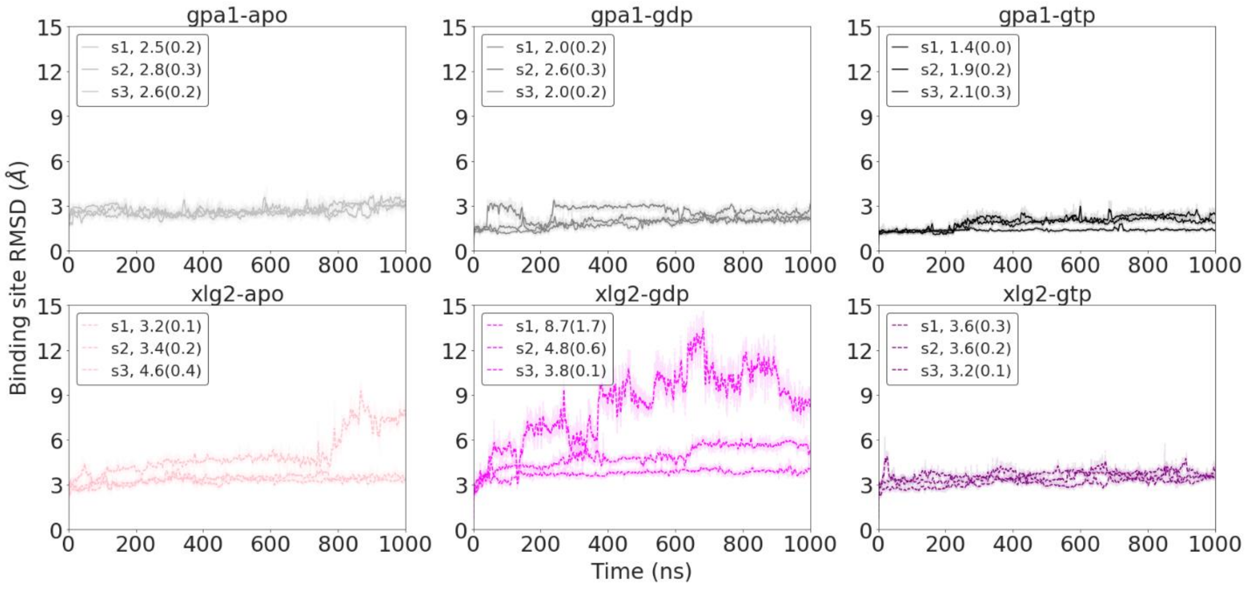

**Figure S6**. Timeseries of the nucleotide binding site RMSD (see Methods for details). The reference structure for GPA1 was the minimized crystal structure, and the minimized homology model for XLG2. The data indicates that the binding site of apo, GDP- and GTP-bound AtGPA1 remains stable throughout the simulations with low deviations. The binding site of XLG2-GTP is stabilized in a relatively stable conformation, whereas the binding site of the GDP-bound state is more unstable and obtains several conformationally different states. Apo XLG2 binding site remains stable for majority of the simulations, which explains its ability to bind AtRGS1 as measured in our experimental studies (see the main manuscript for details).

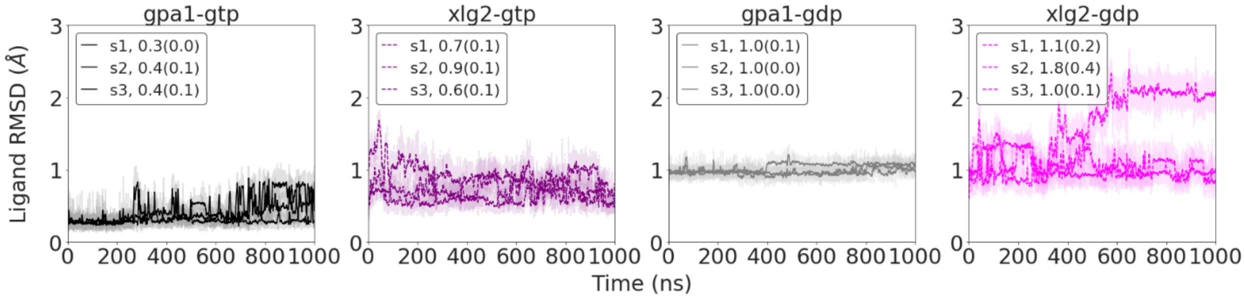


**Figure S7.** Timeseries of ligand RMSD. The data shows that GTP tends to be less mobile than GDP throughout the simulations. And, while GDP is relatively stable in AtGPA1 it is significantly more mobile when bound to XLG2. Interactions between the ligand and protein as well as intra-protein interactions within the binding site maintaining its shape as described in the main manuscript are the key contributors to this behavior of the ligand.

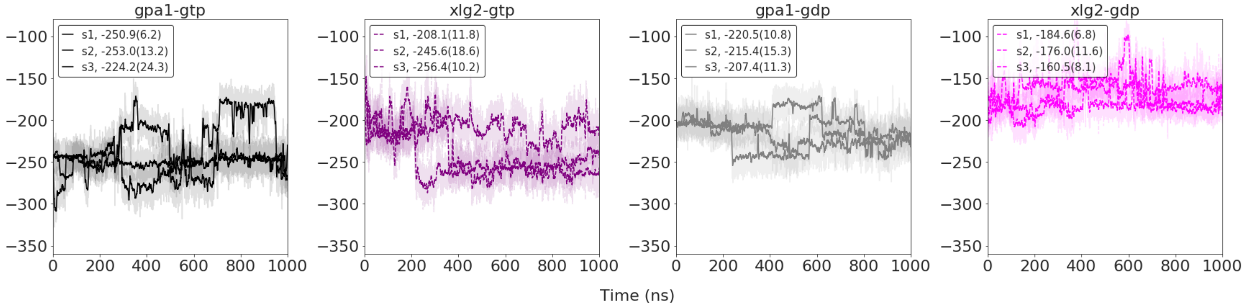

**Figure S8**. Timeseries of the nucleotide-protein interaction energies computed as the sum of Coulomb and LJ potentials in kcal/mol. The trend of the interaction energies are equivalent to the mobility of the ligand in the binding site (Fig. S7), which in turn is contributed by the ligand-protein as well intra-protein interactions maintaining the shape of the binding site (see the main manuscript for details).

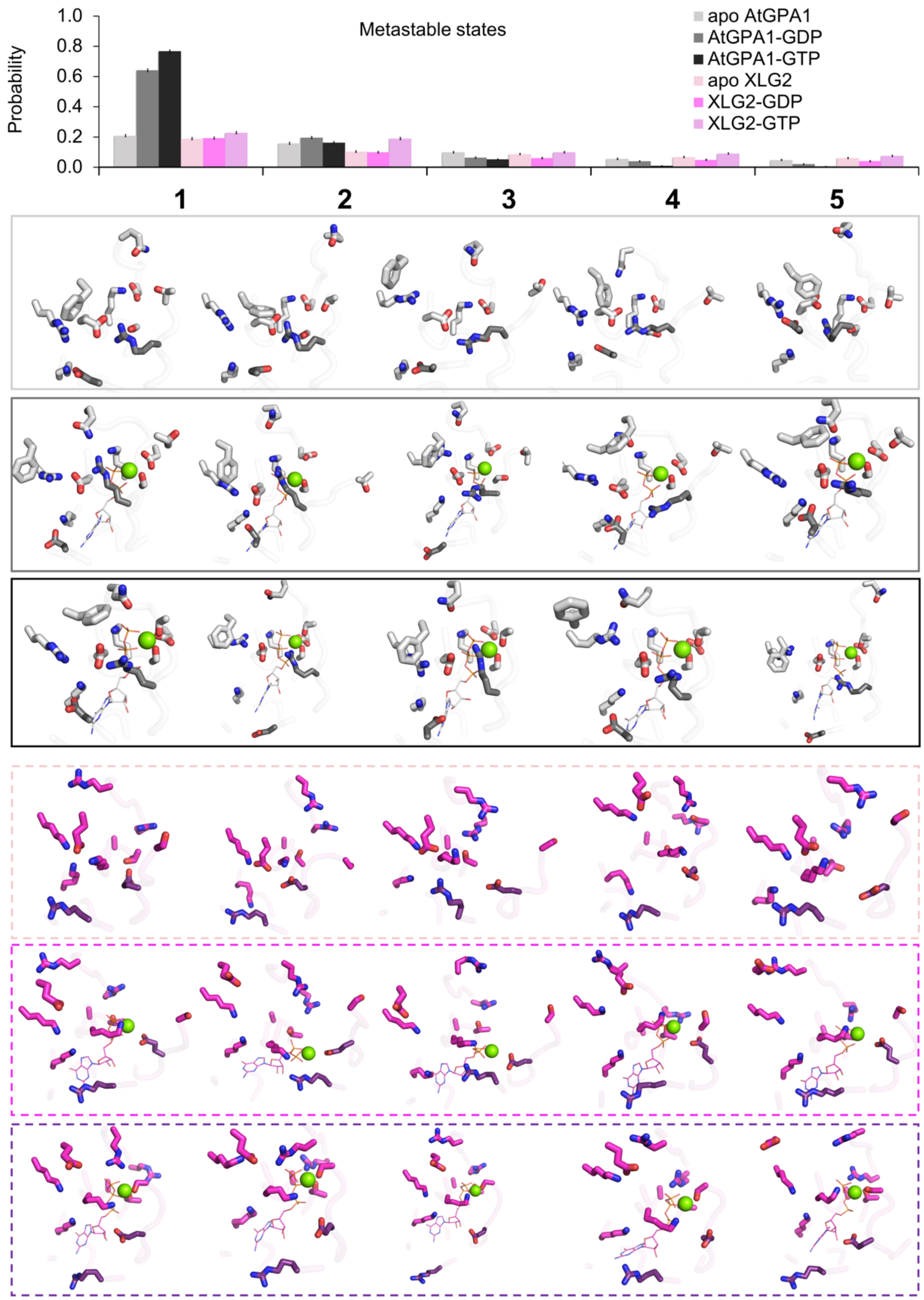


**Figure S9**. Centroid conformations of the top five most populated clusters. The bar plot of the state probabilities shown on top of the figure is the same as in Fig. 6A, and is shown here for convenience. The top five most populated metastable states of the nucleotide binding site obtained in cluster analysis (see Methods for details) indicate that nucleotide-bound AtGPA1 obtains a stable frequently visited and dominant conformational states, whereas XLG2 complexes tend to transition between conformationally diverse states with lower probabilities. Both apo proteins obtain multiple states with equivalently low probabilities.

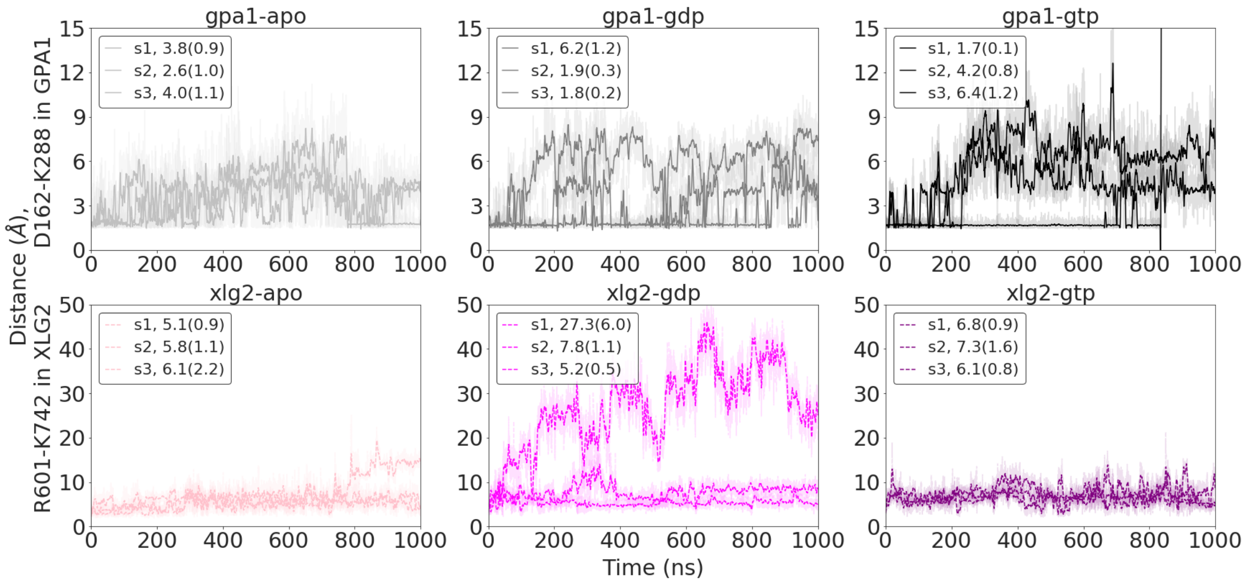


**Figure S10**. Timeseries of D162-K288 distance in GPA1 and the distance between equivalently positioned residues in XLG2 homology model, R601-K742. These residues shape the nucleotide binding sites and impact the nucleotide binding (see the main manuscript for details).

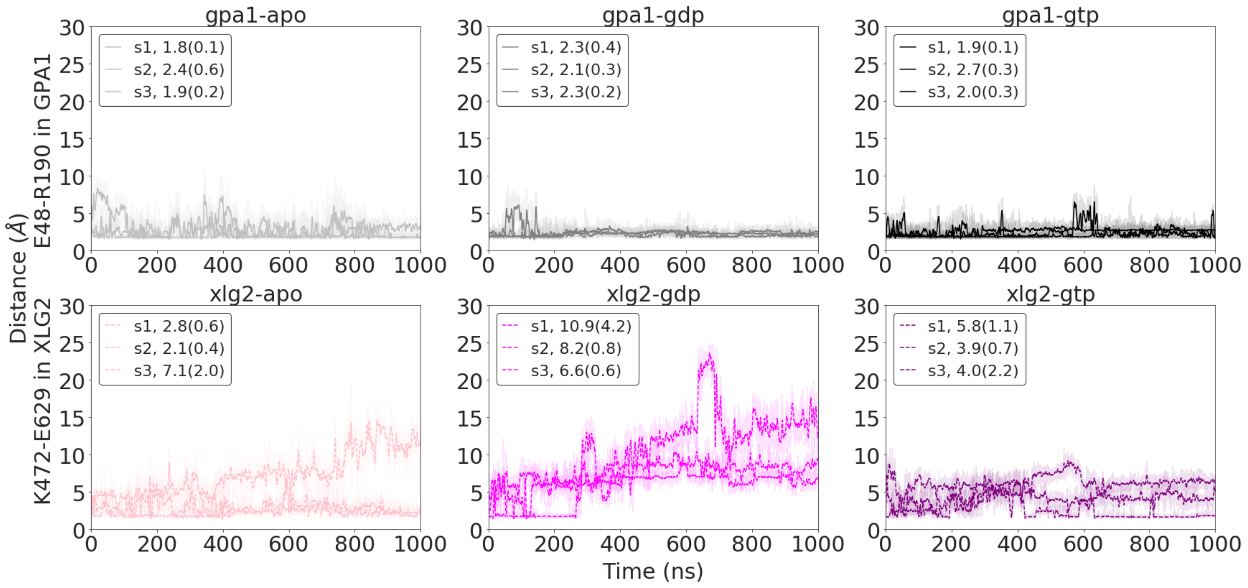


**Figure S11**. Timeseries of E48-R190 distance in GPA1 and K472-E629 distance in XLG2. These residues form the nucleotide binding sites and impact the nucleotide binding (see the main manuscript for details).

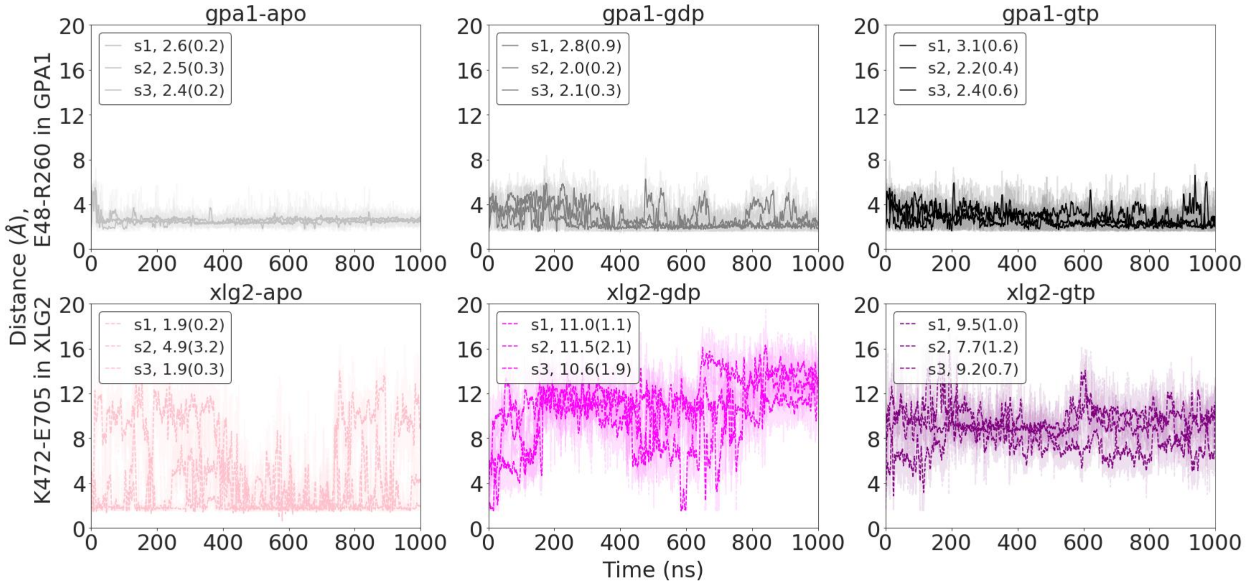


**Figure S12**. Timeseries of E48-R260 distance in GPA1 and K472-E705 distance in XLG2. These residues shape the nucleotide binding sites and impact the nucleotide binding (see the main manuscript for details).

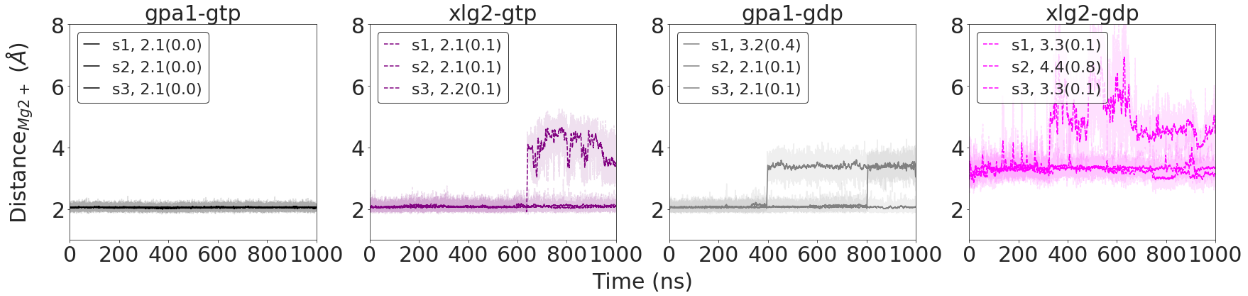


Figure S13. Timeseries of the minimum distance between three $Mg^{2+}$-binding residues, S52, T193, D218, and $Mg^{2+}$ in GPA1, and three $Mg^{2+}$-binding residues, T476, S632, R669, and $Mg^{2+}$ in XLG2. These residues shape the nucleotide binding sites and impact the nucleotide binding (see the main manuscript for details).

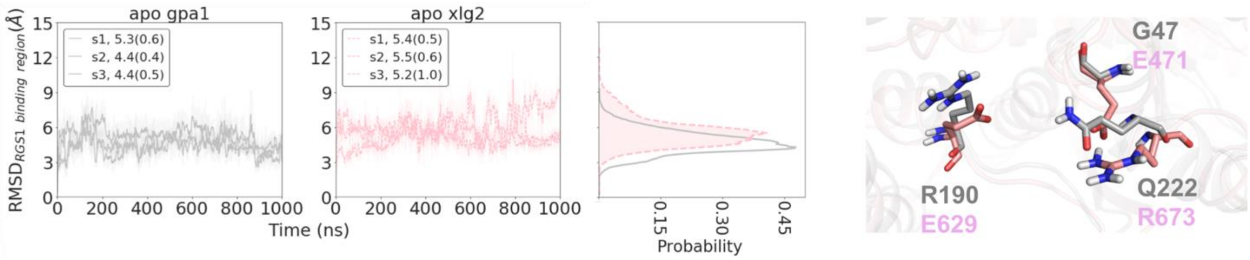


**Figure S14**. Timeseries and distributions of RMSD of AtRGS1 binding site in apo AtGPA1 (light grey) and apo XLG2 (pink). The closely overlapping distributions indicate overall similar structural stability of the regions providing a mechanistic characterization of our experimental results showing equivalent binding by the two proteins to AtRGS1 (see the main manuscript for details).

**Supplementary movie S1.** First five most populated clusters of apo GPA1 in the order from most to least populated. The following residues were used for clustering and are shown in in sticks representation: E48, K51, S52, D162, R190, T193, D218, Q222, F253, R260, K288. The lengths of the states are proportional to the state probabilities.

**Supplementary movie S2.** First five most populated clusters of the GPA1-GDP complex in the order from most to least populated. The following residues were used for clustering and are shown in in sticks representation: E48, K51, S52, D162, R190, T193, D218, Q222, F253, R260, K288. The lengths of the states are proportional to the state probabilities.

**Supplementary movie S3.** First five most populated clusters of the GPA1-GTP complex in the order from most to least populated. The following residues were used for clustering and are shown in in sticks representation: E48, K51, S52, D162, R190, T193, D218, Q222, F253, R260, K288. The lengths of the states are proportional to the state probabilities. The states with populations smaller than 1% are not represented in the movie (states 4 and 5).

**Supplementary movie S4.** First five most populated clusters of apo XLG2 in the order from most to least populated. The following residues were used for clustering and are shown in in sticks representation: K472, A475, T476, R601, E629, S632, R669, R673, E705, K714, K742. The lengths of the states are proportional to the state probabilities.

**Supplementary movie S5.** First five most populated clusters of the XLG2-GDP complex in the order from most to least populated. The following residues were used for clustering and are shown in in sticks representation: K472, A475, T476, R601, E629, S632, R669, R673, E705, K714, K742. The lengths of the states are proportional to the state probabilities.

**Supplementary movie S6.** First five most populated clusters of the XLG2-GTP complex in the order from most to least populated. The following residues were used for clustering and are shown in in sticks representation: K472, A475, T476, R601, E629, S632, R669, R673, E705, K714, K742. The lengths of the states are proportional to the state probabilities.
